# Structural basis for Lamassu-based antiviral immunity and its evolution from DNA repair machinery

**DOI:** 10.1101/2025.04.02.646746

**Authors:** Matthieu Haudiquet, Arpita Chakravarti, Zhiying Zhang, Josephine L. Ramirez, Alba Herrero del Valle, Paul Dominic B. Olinares, Rachel Lavenir, Massilia Aït Ahmed, M. Jason de la Cruz, Brian T. Chait, Samuel H. Sternberg, Aude Bernheim, Dinshaw Patel

**Affiliations:** Institut Pasteur, Université Paris-Cité, CNRS UMR2535, Molecular Diversity of Microbes, Paris, France; Institut Curie, PSL Research University, INSERM U932, Innate Immunity in Physiology and Cancer Team, Paris, France; Structural Biology Program, Memorial Sloan-Kettering Cancer Center, New York, NY 10065; Department of Biochemistry and Molecular Biophysics, Columbia University, New York, NY, USA; Howard Hughes Medical Institute, Columbia University, New York, NY, USA; Laboratory of Mass Spectrometry and Gaseous Ion Chemistry, Rockefeller University, New York, NY, 10065

## Abstract

Bacterial immune systems exhibit remarkable diversity and modularity, as a consequence of the continuous selective pressures imposed by phage predation. Despite recent mechanistic advances, the evolutionary origins of many antiphage immune systems remain elusive, especially for those that encode homologs of the Structural Maintenance of Chromosomes (SMC) superfamily, which are essential for chromosome maintenance and DNA repair across domains of life. Here, we elucidate the structural basis and evolutionary emergence of Lamassu, a bacterial immune system family featuring diverse effectors but a core conserved SMC-like sensor. Using cryo-EM, we determined structures of the *Vibrio cholerae* Lamassu complex in both apo- and dsDNA-bound states, revealing unexpected stoichiometry and topological architectures. We further demonstrate how Lamassu specifically senses dsDNA *in vitro* and phage replication origins *in vivo*, thereby triggering the formation of LmuA tetramers that activate the Cap4 nuclease domain. Our findings reveal that Lamassu evolved via exaptation of the bacterial Rad50-Mre11 DNA repair system to form a compact, modular sensor for viral replication, exemplifying how cellular machinery can be co-opted for novel immune functions.

## INTRODUCTION

Defense systems are highly diverse, reflecting the extensive and ongoing phage-bacteria coevolution ^1,2^. While molecular mechanisms of bacterial immune systems are being rapidly deciphered, the emergence and evolution of these immune systems have remained elusive. Evolutionary studies have until now focused on CRISPR-Cas systems, uncovering their evolutionary origins in transposable elements machinery or toxin-antitoxin systems ^3,4^, but the evolutionary origin of the hundreds of other identified antiphage systems remain to be investigated.

Structural Maintenance of Chromosome (SMC) proteins are a conserved family of ATPases critical for DNA processing across all domains of life ^5,6^. They mediate distinct processes, such as condensation and organization of chromatin, DNA repair, and innate immunity ^7–9^. A notable, widespread, and highly diverse defense system encoding an SMC-like core component is the Lamassu family ^7–9^. It comprises LmuB, an SMC-like protein, LmuC, a small protein with a domain of unknown function, and LmuA, which encodes a range of diverse effectors that include Cap4 nucleases. The extensive diversity of LmuA effectors contributes to the functional versatility of the Lamassu family. Lamassu from the El Tor *Vibrio cholerae* strain (also known as DNA defense module DdmABC), the most studied Lamassu homolog, is thought to be triggered by palindromic sequences from phages and plasmids leading to cell death, thus protecting the bacterial population from further infection ^10,11^.

From a structural perspective, a number of structural modeling and domain analysis of the Lamassu components have revealed the presence of SMC scaffold for the LmuB subunit ^11^, whose coiled coils are approximately half of the length (∼20 nm) of conventional SMC proteins ^7^, such as condensins ^12^, cohesins ^13–15^, Smc5/6 ^16^, MukBEF ^17,18^ and Wadjet JetC ^19^. This non-canonical feature of the Lamassu complex might play a key role in its immune activity. The functional implications of this shorter coiled-coil and its influence on Lamassu complex assembly and specificity remain unclear. How Lamassu complexes detect foreign DNA, trigger effector activation, and evolved their modular architecture remains unknown—posing fundamental questions about the structural and evolutionary principles underlying this widespread antiphage system.

Here we set out to understand the structural basis of Lamassu complex formation and activation and how these structural features illuminate Lamassu evolution across species.

## RESULTS

### Short Lamassus are highly diverse antiphage systems with co-evolving components

We first set out to comprehensively identify Lamassu systems in prokaryotic genomes. We assembled a dataset incorporating experimentally validated Lamassu systems from previous studies ^7,9–11^ as well as predicted systems ^8,9^. We identified Lamassu systems through an iterative approach, aiming to uncover the complete diversity of the system (see Methods, **Fig. 1a**). This approach yielded 3,829 Lamassu systems across 22,920 complete genomes from the RefSeq database, 12% (n=430) more than previous detections on the same genomes. We found Lamassu systems in 3,544 genomes, including seven archaeal genomes (*Euryarchaeota* and *Thaumarchaeota*) indicating that 15.4% of prokaryotic genomes encode at least one Lamassu system, with some species like *Photorhabdus laumondii* encoding up to five distinct Lamassu systems.

**Figure 1:**
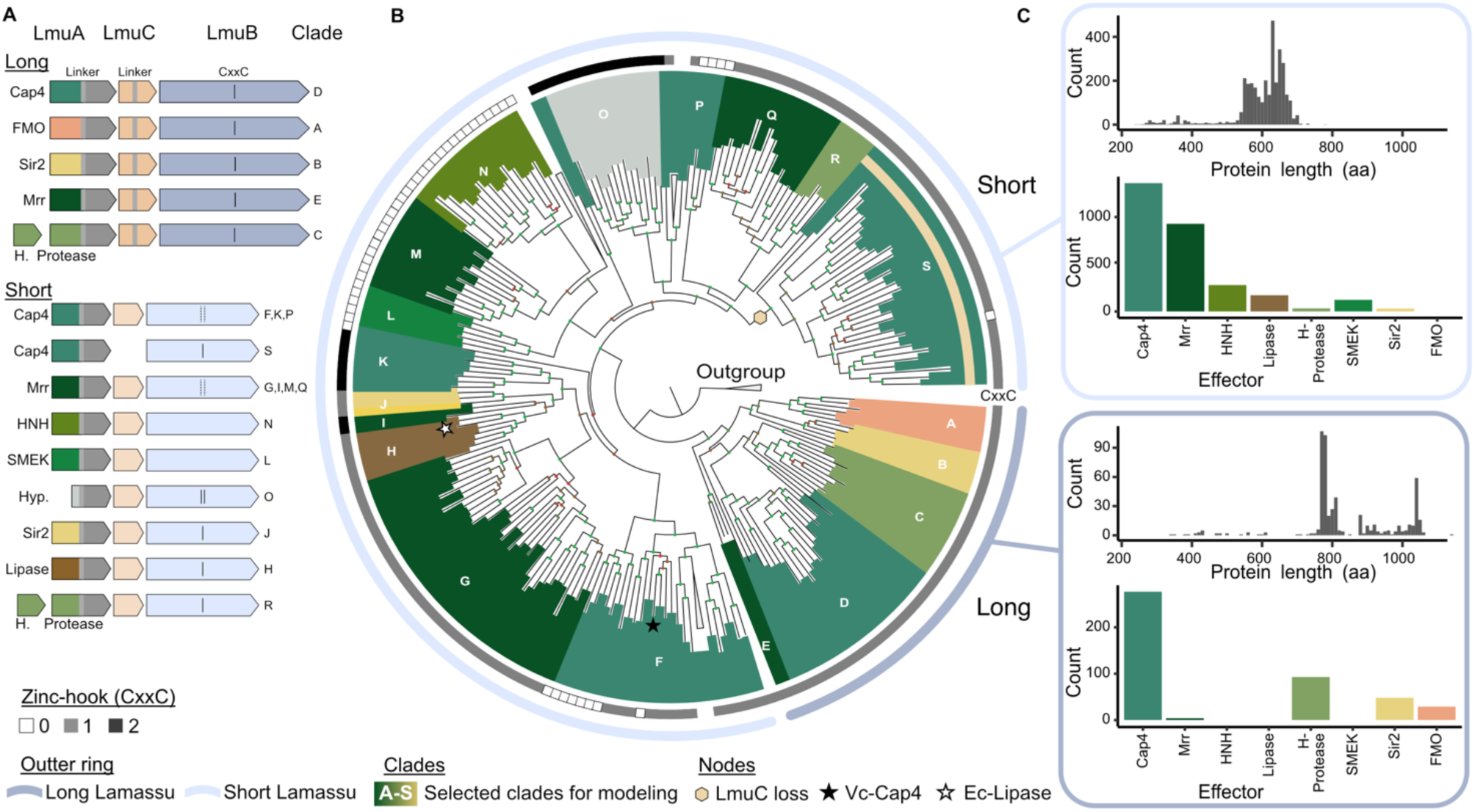
Short Lamassus are highly diverse and modular antiphage systems. **(a)** Types and subtypes of Lamassu identified in DefenseFinder. Representative operons illustrate modularity, highlighting LmuA effector domains (colored), linker (light grey), and C-terminal domain (dark grey). LmuC variants are shown with (long) or without (short) linkers. LmuB variants feature zinc-hook motifs (CxxC); solid black lines indicate all members have the motif, dotted lines indicate presence only in some members. Clade labels correspond to those in panel B. **(b)** Phylogeny of LmuB proteins based on whole-protein alignments of representative sequences clustered at 40% identity and coverage (n = 270). Inner colors at branch tips correspond to LmuA effector domains as in panel A. The beige hexagon indicates hypothesized loss of LmuC in clade S; black star marks the studied Lamassu Vc-Cap4 Lamassu (ddmABC), and white star the Lamassu Ec-Lipase system. The first ring indicates zinc-hook motif copy number; outer ring distinguishes short (light blue) from long (dark blue) types. **(c)** Distribution of LmuB protein lengths and LmuA effector domains across Lamassu systems.

To gain a deeper understanding of the diversity of the family, we focused on LmuB, as it is conserved across all Lamassu systems. Phylogenetic analysis of LmuB uncovered two major clades, which were mainly distinguished by the length of coiled-coil regions: short LmuB proteins averaged 600 amino acids (total protein length), whereas long LmuB proteins measured ∼800 amino acids (**Fig. 1b,c**). 3D structure comparison of LmuB AlphaFold 3 (AF3) models also corroborated this dichotomy (**Extended Data Fig. S1**), and both types were associated with defensive environments, with defense scores of 27% (long) and 25% (short; see Methods). Long and short Lamassus differed in abundance, with 3,293 systems classified as short and 536 as long. Previous experimentally validated systems belong to the short clade^10,11,20^, except one system from *Bacillus cereus* B4077, encoding Hydrolase-Protease effectors ^9^. Long and short LmuB were associated with different LmuA and LmuC gene families, suggesting that the dichotomy extends to LmuB’s partners. Notably, LmuC and the C-terminal domain (CTD) of LmuA that associated with long LmuB were also longer than the ones associated with short LmuB, which was also evident from a 3D structure comparison (**Extended Data Fig. S1**). These findings further underscore the presence of two distinct Lamassu families

We next examined the conservation and diversity of Lamassu features between long and short Lamassus components. LmuB from both types displayed a prototypical Walker A (Gx4GK[S/T]), but their Walker B motifs differed: long Lamassu had the consensus sequence hhhhDD, whereas short Lamassu featured hhhhD[Q/S/T/E] (**Extended Data Fig. S2**). This variation may modulate the activity of the ATPase, for example E→Q mutants can result in efficient ATP binding but decreased ATP hydrolysis ^21^. The putative dimerization module of LmuB, the CxxC zinc-hook, exhibited extensive variation across the LmuB phylogeny. Long LmuB consistently encoded a single copy, while short LmuB encoded either zero, one, or two copies (**Fig. 1a,b, Extended Data Fig. S2**). This suggests that short LmuB may employ divergent dimerization strategies compared to other SMC-like proteins.

In terms of effectors, Lamassu is exceptionally diverse, with at least nine distinct domains found in the N-terminal region of LmuA (**Fig. 1a**). The repertoire of effectors is bigger in LmuAs associated with short (n=8) Lamassus when compared to long ones (n=4). Some effectors are shared between the two types, including Cap4, Mrr, Protease (accompanied by a hydrolase-like protein), and Sir2, while others are type-specific like the flavin-containing monooxygenase (FMO) for long Lamassu, and HNH, SMEK, and Lipase for short Lamassu. To determine whether the shared effector domains—Cap4, Mrr, Protease, and Sir2— arose from the same ancestral domain between long and short Lamassu systems, we analyzed their phylogenies within the broader context of their homologs (**Extended Data Fig. S3**). In all cases except for Sir2, effectors from long and short Lamassu formed distinct but closely related sister clades, suggesting either vertical inheritance or exchange between Lamassu systems. In contrast, Sir2 domains exhibited a markedly divergent evolutionary trajectory, with long- and short-associated Sir2 homologs branching apart in the phylogeny. This pattern suggests that while Cap4, Mrr, and Protease were likely inherited or transferred between Lamassu systems, Sir2 effectors originated from independent acquisition events. Finally, the composition of Lamassu varied significantly in the short Lamassu families. We observed loss of LmuC in a specific clade (Clade S, **Fig. 1b**), an architecture previously called type I Lamassu shown to be antiphage ^7^. We also identified a clade of short Lamassu which seemingly lost the effector domain of LmuA (Clade O, **Fig. 1b**). This clade contained more than 100 sequences across more than 40 species and was associated with non-truncated LmuC and LmuB genes.

Overall, short Lamassus are more diverse in terms of sequence, effectors and more abundant in bacterial genomes. We therefore focused on short Lamassus, selecting the experimentally validated Cap4 system from *Vibrio cholerae* (Lamassu Vc-Cap4) for structural elucidation. This system encodes an LmuB with no zinc-hook, an LmuC, and a Cap4-like LmuA effector predicted to degrade double-stranded DNA (**Fig. 2A**).

**Figure 2:**
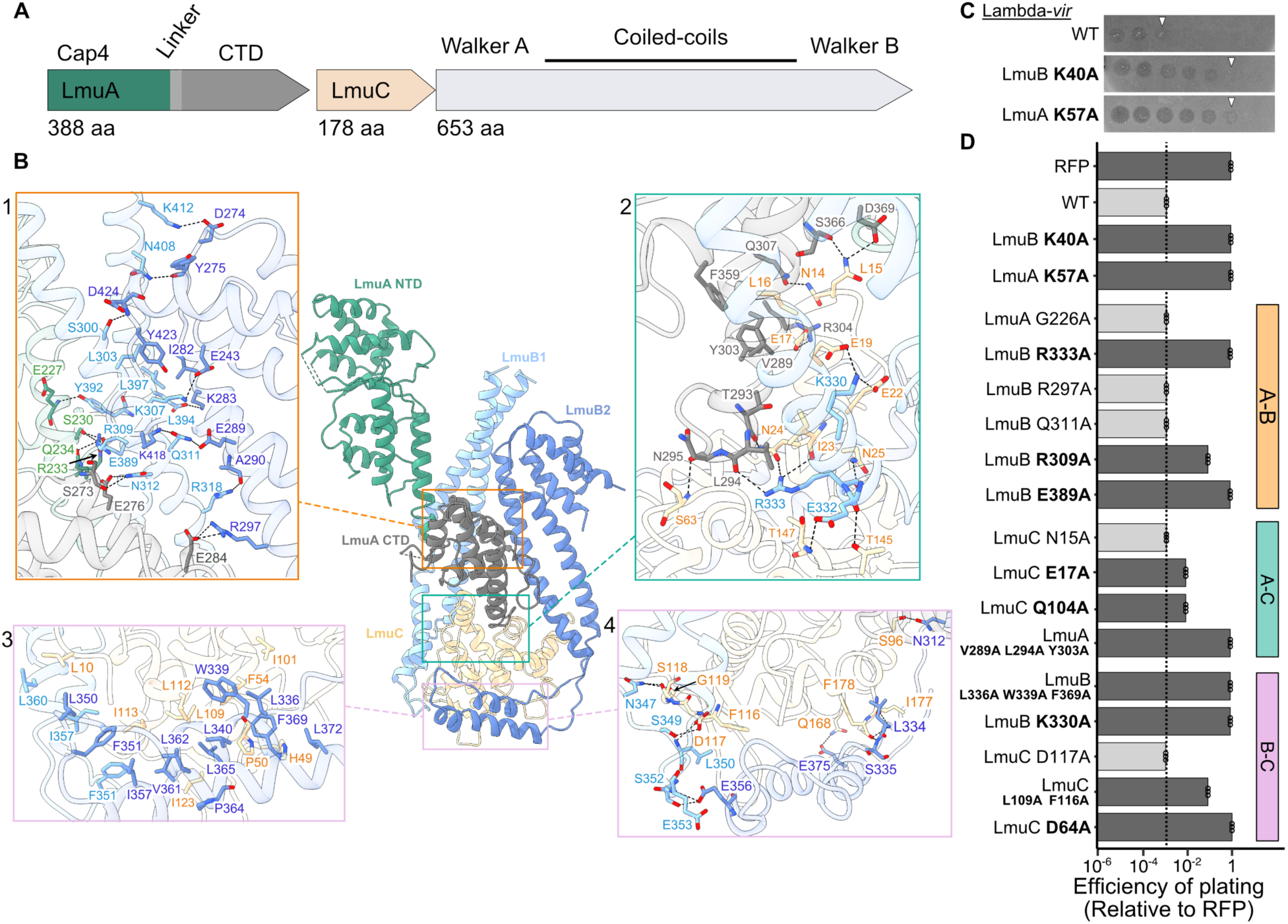
Apo-state complex of LmuA_1_B_2_C_1_. **(a)** Operon schematic of Lamassu Vc-Cap4 using color coding from Fig. 1a. **(b)** Cryo-EM structure of the apo-LmuA_1_B_2_C_1_ complex shows core interactions involving LmuB (light/dark blue), LmuC (beige), and LmuA CTD (dark grey), with minimal interactions involving LmuA’s N-terminal Cap4 domain (green). Insets (1–4) illustrate detailed hydrogen bonding and hydrophobic interactions, also shown in Extended Data Fig. S7. **(c)** Representative plaque assays of WT Lamassu Vc-Cap4 and catalytic mutants (LmuB K40A, LmuA K57A) against Lambda-vir phage. **(D)** Plaque assays probing structural interfaces. Efficiency of plating (EOP) indicates the relative plaque-forming efficiency of the tested phage on the bacterial strain compared to the negative control strain (RFP). An EOP of 1 indicates a completely non-functional defense system (no restriction of phage replication). Mutants impairing activity (dark grey bars) are highlighted in bold; WT activity level indicated by dotted line. Data points represent three biological replicates. Targeted interfaces (A-B, A-C, B-C) are shown on the right.

### Apo-LmuABC complex reveals an unanticipated core topology that sequesters monomeric LmuA

Our initial attempts to purify LmuABC from the full operon (**Fig. 2a**) were unsuccessful, prompting us to express its components separately. We were able to co-express and purify the LmuBC subcomplex, however separately purified LmuA exhibited severe aggregation upon size-exclusion chromatography (SEC) analysis (**Extended Data Fig. S4a**). For LmuBC, SEC yielded a well-defined peak containing both LmuB and LmuC, which migrated at their expected sizes on SDS-PAGE (**Extended Data Fig. S4b**). Notably, this copurification of tagged LmuB coeluting with LmuC, indicated a stable interaction between the two proteins. To reconstitute the full LmuABC complex, we mixed purified LmuA and LmuBC from their respective heparin columns followed by running it on a size exclusion column. SEC (**Extended Data Fig. S4c**) and native mass spectrometry (nMS) (**Extended Data Fig. S5a**) confirmed the formation of apo-LmuA₁B₂C₁. Addition of 20-bp DNA resulted in formation of an LmuA₁B₂C₁-dsDNA complex (**Extended Data Fig. S5b**).

Cryo-EM analysis of the LmuABC complex in the presence of a 20-bp dsDNA substrate revealed the presence of both the apo and dsDNA-bound states. To confirm the identity of the apo complex, we independently obtained the structure of the LmuABC complex in the absence of dsDNA, yielding an identical assembly. As both apo-LmuABC structures showed identical conformations and assemblies, with the apo-one obtained from the complex with dsDNA having higher resolution (3.21 Å), we decided to focus on this structure for further analysis (**Fig. 2a,b**; work-flows in **Extended Data Fig. S6** and data statistics in **Extended Data Table 1**). This apo-LmuABC structure revealed a single full-length LmuA, the head-distal coiled coils of two LmuB subunits adopting different interconnected folds, and one full-length LmuC, with these interacting segments forming the core of the LmuABC complex. The kinked head-distal coiled coils of LmuB₁ (single kink) and LmuB₂ (a pair of kinks) converge at the distal segment, interacting with the CTD of LmuA and the globular fold of centrally positioned LmuC. Notably, LmuC directly associates with these coiled coils, forming an integral part of the complex (**Fig. 2b, Extended Data Fig. S7**). The Cap4 N-terminal domain of LmuA exhibits minimal interactions with the other subunits. A key feature of the apo-state is the monomeric nature of LmuA, with its NTD in an extended conformation relative to its CTD, with the latter stabilized through interactions with its partner subunits. Since LmuC was previously predicted to function as a kleisin ^8^, its placement within the LmuABC core was unexpected, given that in conventional SMC complexes, kleisins typically associate asymmetrically with the head-proximal coiled coils and head domains ^15^.

To investigate the functional relevance of the apo-LmuABC structure, we performed mutational analysis and assessed Lamassu’s antiphage activity *in vivo* (**Fig. 2c,d**). We expressed the natural operon of Lamassu Vc-Cap4 under an inducible promoter on a medium-copy plasmid in *E. coli*. In this setup, at low induction levels of 0.0001% L-Arabinose, Lamassu is strongly antiviral against Lambda-vir, but not toxic (**Fig. 2c, Extended Data Fig. S9**). In contrast, the Lamassu Vc-Cap4 system is toxic at high inducer concentration, except for LmuA K57A mutant, suggesting auto-activation at high intracellular concentration (**Extended Data Fig. S8**). The LmuB-LmuC (B-C) interface was targeted through alanine substitutions at K330A in LmuB, and D64 and D117 in LmuC (**Fig. 2b,d**, **Extended Data Fig. S7**). Strikingly, all mutations except D117A abolished antiphage activity, highlighting the critical role of the B-C interface in system function (**Fig 2d**). This was further supported by disrupting hydrophobic patches at L336+W339+F369 in LmuB and L109+F116 in LmuC, both of which completely abolished activity (**Fig. 2b,d**, **Extended Data Fig. S7**). Similarly, targeting the LmuA-LmuB (A-B) interface—R333A, R309A and E389A in LmuB—abolished antiphage activity, reinforcing the functional significance of these interactions (**Fig. 2b,d**, **Extended Data Fig. S7**). For the LmuA-LmuC (A-C) interface, alanine mutations were introduced at N15, E17 and Q104 in LmuC, as well as V289+L294+Y303 in LmuA (**Fig. 2b,d**, **Extended Data Fig. S7**). Among these, E17A displayed an intermediate phenotype, the hydrophobic patch mutant (V289+L294+Y303 in LmuA) completely disrupted function, whereas the other mutations had no significant effect (**Fig. 2d**). Together, these findings establish the structural and functional importance of the B-C, A-B and A-C interfaces in LmuABC-mediated antiphage defense, while also revealing a previously uncharacterized role for LmuC in the complex architecture.

### Lamassu senses DNA ends *in vitro* and phage origin of replication *in vivo*

To understand how Lamassu effector activity is triggered, we next focused on the dsDNA-bound structure. We determined a 2.93 Å resolution structure, which revealed a complex composed of LmuA₁B₂C₁ bound to a single 20-bp dsDNA molecule (**Fig. 3a,b**, work-flow in **Extended Data Fig. S6** and data statistics in **Extended Data Table 1**). The structure shows that Lamassu specifically binds the dsDNA end, rather than engaging along its length as might have been expected from other SMC complexes. Structural alignment of the apo- and dsDNA-bound complexes resulted in an RMSD of 0.795 Å (**Extended Data Fig. S9**), indicating that while the global architecture is largely maintained, DNA binding induces localized rearrangements, particularly in LmuB₁, where additional density allowed tracing of its head domain and the head-proximal coiled coil that were too flexible to be resolved in the apo-state. By contrast, the head domain and head-proximal coiled coil of LmuB₂ remain flexible, suggesting asymmetry in DNA engagement.

**Figure 3:**
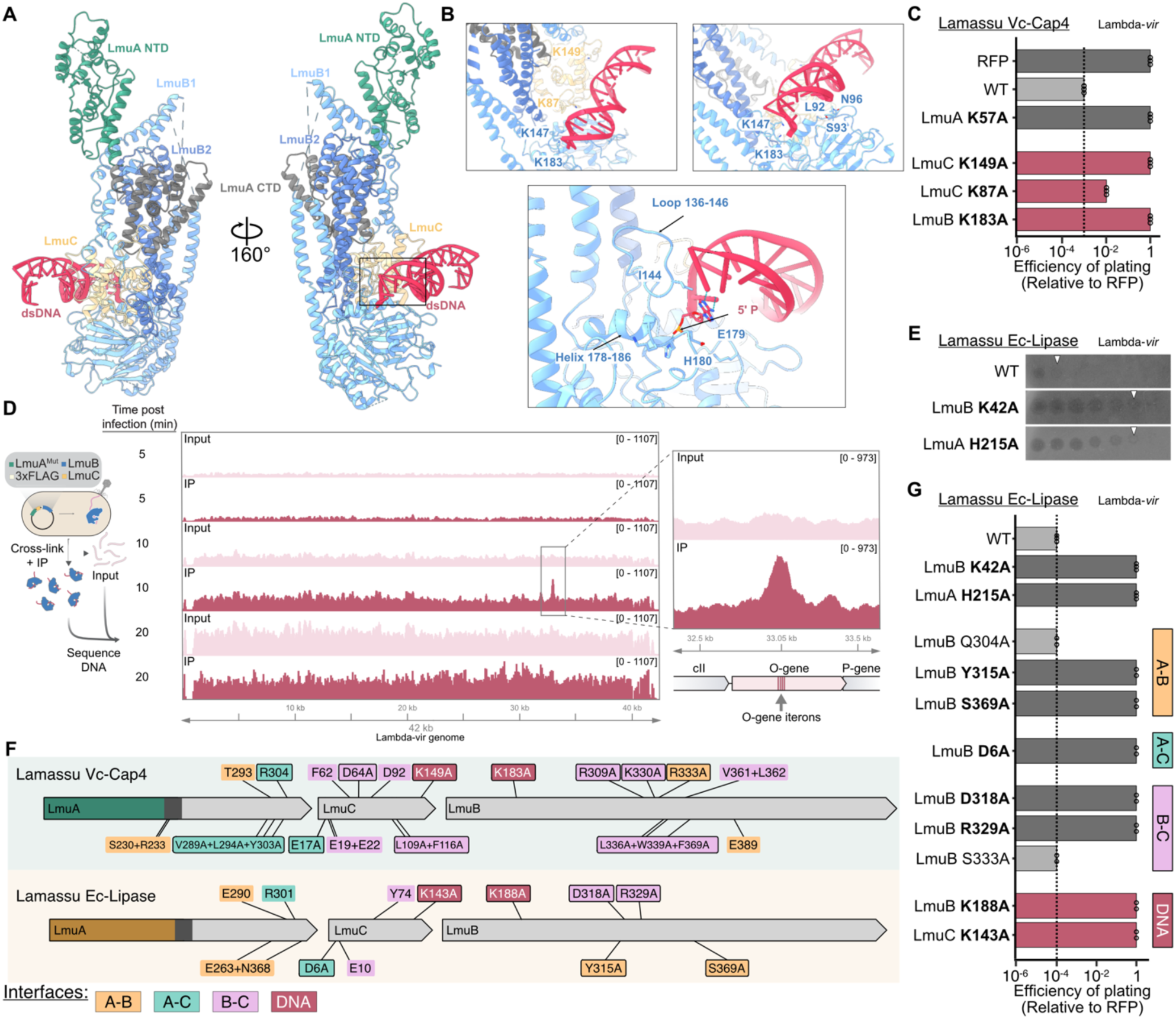
Lamassu binds DNA ends *in vitro* and phage origins of replication *in vivo*. **(a-b**) Cryo-EM structure of LmuA_1_(K57A)B_2_C_1_ bound to 20-bp dsDNA. Insets highlight specific protein-DNA interactions that block DNA trajectory. **(c)** Plaque assays demonstrating essential roles of residues LmuC K149 and LmuB K141 in antiphage defense, with partial impairment observed for LmuC K87A. **(d)** Schematic of ChIP-seq experimental workflow with FLAG-tagged LmuB, LmuC, and catalytically inactive LmuA (K57A Mut; left), and representative ChIP-seq data (middle) from experiments in which cells were infected with Lambda-vir. The genome-wide graphs show normalized read coverage mapped to the Lambda-vir genome at various time points after infection, with input and IP samples colored in pink and maroon, respectively. FLAG-tagged LmuB specifically enriched the Lambda-vir origin (O gene iterons) 10 min after infection, as shown in the magnified inset at right; the genomic locus is annotated below the graph. **(e)** Validation of Lamassu Ec-Lipase antiphage activity by plaque assay. **(f)** Comparison of interface residues between Lamassu Vc-Cap4 and Lamassu Ec-Lipase; residues critical for defense verified by assays are indicated and shown in Fig. 2d, and Fig. 3g. **(g)** Plaque assay of mutants of Lamassu Ec-Lipase. Efficiency of plating indicates the relative plaque-forming efficiency of the tested phage on the bacterial strain compared to the negative control strain (RFP). An EOP of 1 indicates a completely non-functional defense system. Interfaces probed are indicated on the right; mutants with impaired activity (dark grey bars) indicated in bold. WT activity level shown by dotted line. Data points represent three biological replicates.

The dsDNA duplex is captured within the LmuABC complex in a conformation that obstructs its path, effectively blocking further progression through the structure. Residues contributing to this obstruction include I144, located in a loop spanning residues 136–149, and E179 and H180, positioned on an α-helix spanning residues 178–186. In this helix, LmuB H180 contacts the dsDNA end at the 5’ phosphate (**Fig. 3b**). These elements appear to define a DNA end binding pocket rather than a DNA-threading mechanism. The DNA interacts primarily with the head domain of LmuB₁ and the globular domain of LmuC. Key residues from the DNA contacting interface include S93, N95, K147, and K183 in LmuB₁, as well as K87 and K149 in LmuC. Unlike canonical SMC complexes, where the head-proximal coiled coils engage dsDNA, the LmuABC complex replaces these interactions with LmuC-DNA contacts.

To determine the functional importance of these residues, we generated LmuB K183A and LmuC K87A and K149A mutants and tested their antiphage activity using a Lambda-vir plaque assay. LmuB K183A and LmuC K149A mutants abolished defense, while LmuC K87A mutant caused a partial reduction in activity (**Fig. 3c**). These results indicate that proper DNA recognition is not solely mediated by LmuB but also requires LmuC. Given its direct role in stabilizing dsDNA binding at the complex interface, LmuC likely functions as a specificity factor, ensuring that LmuABC engages dsDNA ends in a regulated manner. Accordingly, AF3 models of LmuC across the phylogeny of short Lamassu repeatedly hit HTH-type transcription factors (helix-turn-helix) with DALI and FoldSeek. These findings suggest that LmuABC binds dsDNA in a structurally constrained mode, with localized rearrangements supporting selective recognition of DNA ends.

Having shown that Lamassu specifically binds dsDNA, particularly sensing the 5’ end, we hypothesized that it might recognize phage-specific DNA substrates during infection. To identify such substrates in vivo, we performed chromatin immunoprecipitation followed by sequencing (ChIP-seq) of Lamassu during infection with phage Lambda-vir (**Fig, 3d, Extended Data Fig. S10a**). We expressed the catalytically inactive LmuABC complex harboring a K57A LmuA mutation under the arabinose-inducible pBAD promoter, and then performed ChIP-seq of FLAG-tagged LmuB at 5, 10 and 20 minutes post-infection, with the goal of enriching specific substrates recognized by the Lamassu system during infection. The resulting data revealed a sharp and specific enrichment of LmuB around the four iterons — short direct DNA repeats — that comprise the Lambda origin of replication, located within the O gene (**Fig. 3D**). The O gene encodes the O protein, which initiates theta replication by binding its iterons and assembling a nucleoprotein complex that recruits the host replication machinery ^22^. LmuB enrichment around the iterons was detected specifically at 10 minutes post-infection, but not at 5 or 20 minutes, suggesting that Lamassu senses a transient replication intermediate substrate early during infection.

To test whether LmuB recognizes the DNA sequence of the O locus itself, independent of phage replication, we repeated ChIP-seq experiments on cells carrying a plasmid encoding only the enriched locus, but lacking Lambda phage. In this context, no enrichment was observed around the iterons (**Extended Data Fig. 10b**), indicating that Lamassu does not bind specifically to the sequence, but rather to a transient DNA structure during phage infection.

### Lamassu interfaces with DNA are conserved across diverse Lamassus

We aimed to determine whether the atypical organization and DNA binding observed in our structural work represents a recent innovation within the clade of short Lamassu Cap4 (Clade F, **Fig. 1b**), which have lost their Zn-hook. We tested this by experimentally validating a distant system from *E. coli*, referred to as Lamassu Ec-Lipase, which shares only 37.5% sequence similarity and belongs to short Lamassus encoding a lipase effector and a Zn-hook (Clade H, **Fig. 1b**). Similar to Lamassu Vc-Cap4, Lamassu Ec-Lipase was toxic at high inducer concentrations (**Extended Data Fig. S8**) but efficiently protected against Lambda-vir at lower, non-toxic levels (**Fig. 3f**); both the LmuB Walker A mutant (K42A) and the lipase catalytic mutant (H215A) impaired antiphage activity, but only the LmuA H215A mutant was non-toxic at high inducer levels (**Fig. 3e, Extended Data Fig. S8**).

Assuming the atypical Lamassu Vc-Cap4 LmuABC molecular complex is conserved across short Lamassus, we extrapolated the interfaces identified in our structural work to the Lamassu Ec-Lipase system using AF3 multimer models with the stoichiometry determined earlier, comparing these to our Lamassu Vc-Cap4 structure and AF3 model (**Extended Data Fig. S11**). We identified four similar interchain interfaces (A-C, B-C, A-B, and DNA-Lamassu) and found nine analogous interactions: three pairs for A-B, three for B-C, one for A-C, and two involving dsDNA (**Fig. 3f,g, Extended Data Fig. S11**). Alanine mutagenesis of Lamassu Ec-Lipase revealed that seven out of nine mutants abolished its activity, affecting at least one key interacting pair for each interface (**Fig. 3g**). These results, in combination with our AF3 models, support that another member of the short Lamassu clade may form a similar molecular complex that interacts with DNA, with conserved interfaces essential for antiphage function. Hence, LmuABC assembles into an atypical SMC-like complex critical for activity across the diversity of short Lamassus.

### Structural basis of LmuA activation

Since we observed that the LmuABC complex binds to dsDNA ends, and LmuB is a predicted ATPase, we wanted to understand how dsDNA and ATP play a role in Lamassu’s activation.

The presence of a Walker A and Walker B domain in LmuB suggest that ATP binding, and potentially hydrolysis, is required for Lamassu to function. Additionally, mutation of the catalytic Lysine (K40A) of the Walker A impairs defense against Lambda-vir ^9,10^. We therefore tested the *in vitro* activity of LmuABC in the presence or absence of ATP. Our initial biochemical assays revealed non-specific DNA degradation when high concentration of LmuABC (>20 nM) was added to pUC19 plasmid DNA without ATP (**Extended Data Fig. S12a**). To prevent cleavage from autoactivation, we decreased the LmuABC concentration until minimal DNA degradation could be observed. Increasing ATP concentrations correlated with enhanced DNA degradation (**Fig. 4a**), while addition of slowly hydrolyzable ATPγS completely inhibited degradation (**Fig. 4a**). This suggests that ATP binding alone may be insufficient for activation and that ATP hydrolysis to ADP may contribute to LmuABC-mediated DNA processing.

**Figure 4:**
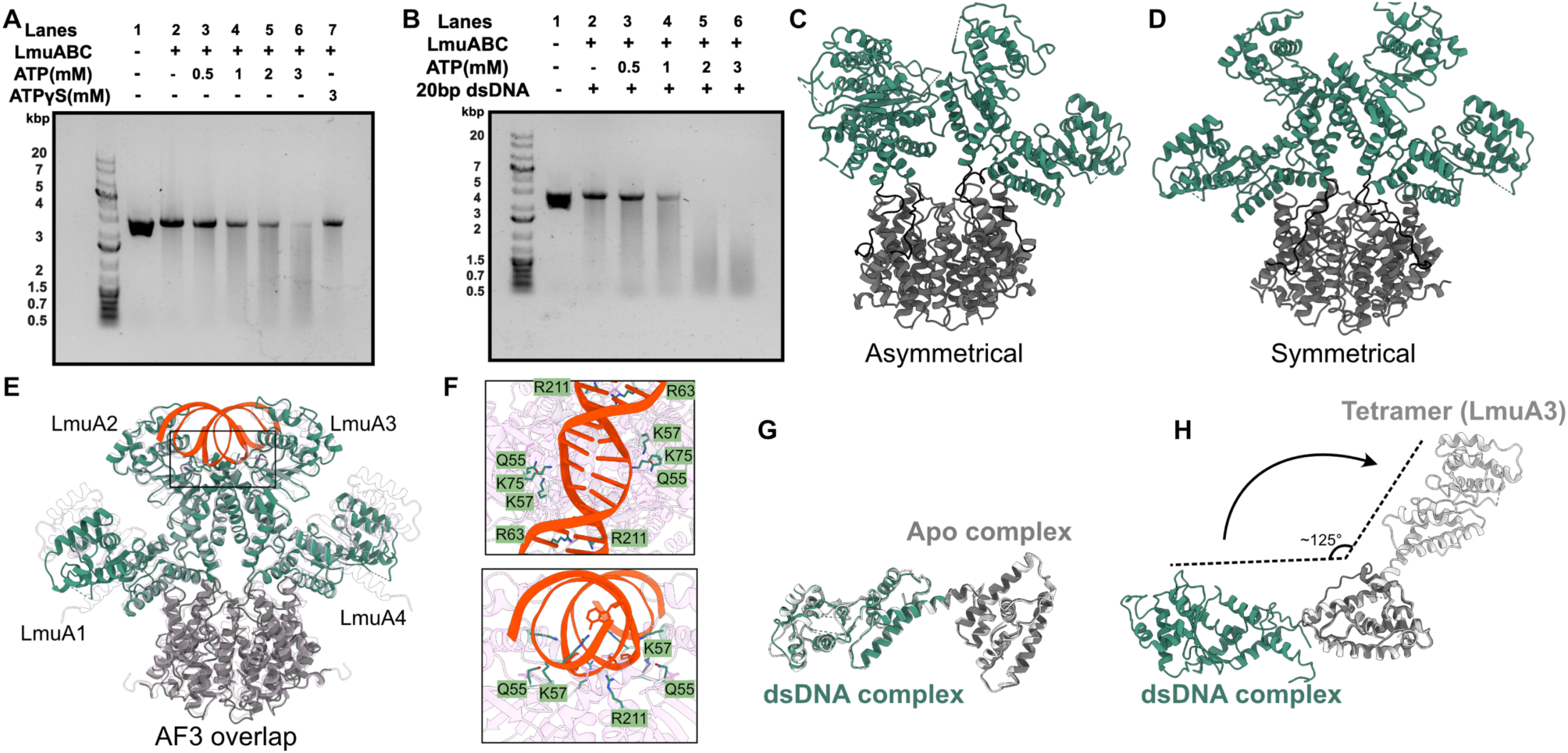
Structural basis of LmuA activation. **(a)** ATP-dependent DNA degradation assay of LmuABC (20 nM) on pUC19 pUC19 (40 nM); increasing ATP concentrations enhance DNA degradation activity of LmuABC. **(b)** Addition of 20-bp dsDNA (0.2 μM) strongly enhances ATP-dependent DNA degradation activity of LmuABC. (**c-d**) Cryo-EM structures of asymmetric (**c**) and symmetric (**d**) LmuA tetramers, illustrating distinct bent and extended conformations stabilized by polar interfaces (shown in Extended Data Fig. S14). (**e**) AlphaFold 3 model (pink) aligned with symmetric tetramer; dsDNA aligns precisely with predicted DNA cleavage site. **(f)** Close-up of residues involved in DNA binding and cleavage predicted by AF3, highlighting catalytic residues including K57. **(g)** LmuA shows no significant conformational changes between apo and DNA-bound complexes (RMSD = 0.795 Å). **(h)** Dramatic conformational rearrangements occur at the linker region between DNA-bound LmuABC and activated LmuA tetramer structures.

Interestingly, DNA degradation was primarily observed at high ATP concentrations in the millimolar range, suggesting that LmuB, which contains a divergent Walker B motif, likely hydrolyzes ATP at a slow rate. Finally, comparison of degradation assays in the absence (**Fig. 4a**) and presence (**Fig. 4b**) of 20-bp dsDNA revealed that addition of this DNA substrate enhanced LmuABC-catalyzed pUC19 degradation. These results suggest that Lamassu’s activity is likely to be regulated both by slow ATP hydrolysis and by interactions with dsDNA that participate in activating the nuclease activity of Cap4.

Our biochemical assays showed that Cap4 activity from LmuA required the presence of LmuABC, and was not detectable with LmuA alone in solution (**Extended Data Fig. S12b**). We hypothesized that DNA end binding by LmuABC could activate LmuA, and sought to capture this active state by cryo-EM. Since the complex is biochemically active at high concentrations (>20nM LmuABC) of LmuABC, we reasoned that adding LmuBC to an excess of LmuA might stabilize active conformations suitable for cryo-EM. Analysis of this sample revealed a tetrameric architecture for LmuA effector—a dimer of dimers—present in both two-fold symmetric and asymmetric conformations at similar abundance. (**Fig. 4c,d, workflow in Extended Data Fig. S13, Table 1)**. LmuA tetramers in both states are held together mainly through interactions involving their C-terminal domains (CTDs) (**Fig. 4c,d**), which superimpose with a RMSD of 2.02 Å **(Extended Data Fig. S14a).** Notably, following superposition of their CTD, the Cap4 NTD of the symmetrical and asymmetrical LmuA conformations display significantly different relative orientations **(Extended Data Fig. S14b)**. Strikingly, the symmetrical tetramer displayed a predicted DNA degradation pocket between LmuA2 and LmuA3 NTDs (**Fig. 4d**).

AF3 modeling using four LmuA monomers and a dsDNA molecule recapitulated the symmetrical tetramer with the dsDNA bound between LmuA2 and LmuA3 (**Fig. 4E**). Hallmark residues comprising K57, which is essential for DNA degradation *in vitro* and phage protection *in vivo*, are aligned in close vicinity of the DNA, forming a DNA cleavage pocket, predicted to result in 1-nt 3’-overhang containing DNA duplexes as the cleavage product (**Fig. 4f**). We thus hypothesized that the LmuA tetramer is the end result of Lamassu activation. Notably, we could not observe significant conformational changes between LmuA in the dsDNA-bound vs. apo-LmuABC complex (**Fig. 4g**), but a significant rotation of the NTD Cap4 domain of LmuA in the active tetramer (**Fig. 4h**). Such a switch in conformation could be facilitated most likely by a complete disengagement of LmuA from the complex upon activation. This disengagement would facilitate the oligomerization into a tetramer, liberating essential residues from the CTD for the tetramerization that are occluded in the apo- or dsDNA-bound complexes and thus allowing the formation of the DNA degradation pocket in the NTD.

### Lamassus evolved from DNA repair complex SbcCD

Lastly, we investigated the likely evolutionary origins of Lamassu. SMC proteins are part of the P-loop NTPase superfamily and have diversified into two major families that can be distinguished by the presence of either a hinge (Hinge SMC) or zinc-hook (CxxC SMC) dimerization domain ^8^. Hinge-encoding SMCs, such as the SMC1–6 complexes in humans and MukB in *Escherichia coli*, are primarily involved in the condensation and organization of chromosomal DNA ^5,23^. CxxC SMCs such as RAD50 in humans and SbcC in *E. coli*, play a critical role in DNA repair, notably of double-strand breaks for RAD50-Mre11-Nsb1^24^ and of palindrome-like convergent replication fork intermediates for SbcCD ^25^. Thus, model CxxC SMCs are primarily implicated in processing non-canonical DNA ends to facilitate subsequent repair steps ^26^. Given the known biochemical activity of Lamassu, and the resemblance of LmuB to SbcC, we thus hypothesized that Lamassu might have emerged from SbcCD.

To test for this, we performed a sensitive search for all homologs of LmuB, resulting in n=367,581 proteins grouped in 1320 families (30% coverage and identity), including proteins from other defense systems such as Rloc ^27^ and DndD, the SMC-like ATPase of the Dnd system ^28^, and numerous SbcC homologs from the SbcD complex but no MukB homolog. This further supports that LmuB is a closer homolog of SMC-like proteins like SbcC than hinge-SMCs like MukB. The resulting phylogeny (**Extended Data Fig. S15**) also recovers the two clades of long and short LmuB. We thus subsampled diverse members of SbcC, long and short Lamassu and the distant SMC protein MukB to root our phylogeny. This revealed that short Lamassu emerged from the clade of long Lamassu, and places long LmuB closer to SbcC (**Fig. 5a**).

**Figure 5:**
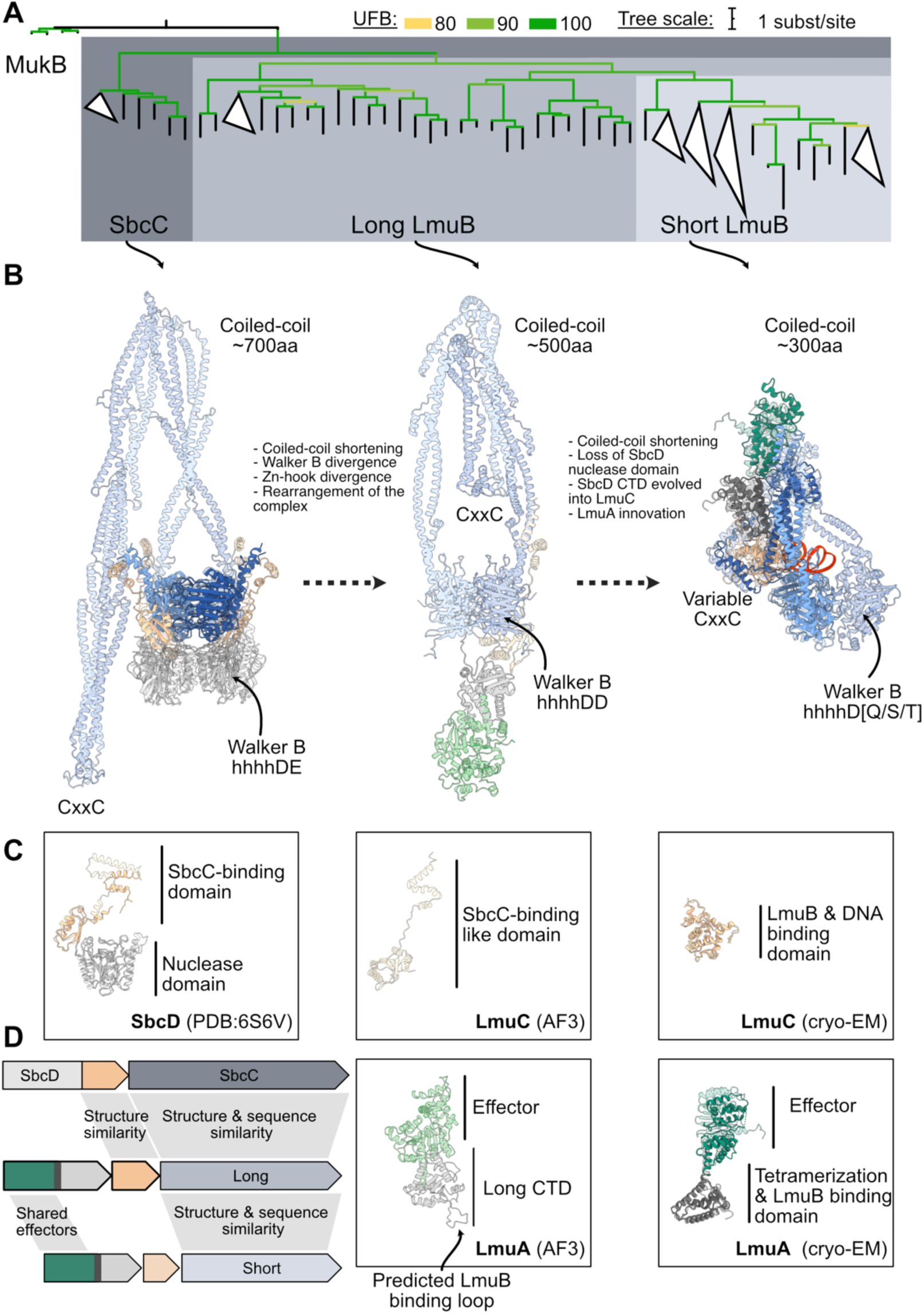
Lamassu originated in SbcCD DNA repair complex. **(a)** LmuB homologs phylogenetic tree. Structure-guided alignment (MukB, SbcC, long LmuB and short LmuB) of AF3 models used as seed alignment for Mafft. The tree was generated with Iq-tree2 using ModelFinder (Q.pfam+F+I+R7) and Ultra-fast Bootstraps (UFB). Nodes with UFB<80 are collapsed. (**b)**. Evolutionary scenario of the emergence of Lamassu from the DNA repair system SbcCD. Experimentally solved structures shown in solid colors, AlphaFold3-modeled structures shown transparently. SbcCD partial complex is taken from PDB:6S7V, and the short Lamassu structures are taken from the cryo-EM structures of DNA-bound Lamassu Vc-Cap4. **(c)** Structural comparison of monomers SbcD, LmuA, and LmuC **(d)** Comparison of typical operons from SbcCD and Lamassu systems highlighting regions of homology (grey shading).

We then studied the evolutionary links of LmuA and LmuC. For LmuC, a similar approach than for LmuB did not recover hits outside the Lamassu system, and overall the size and sequence diversity of LmuC precluded us from relying on sequence alignment methods. Using structural modeling and structure-similarity search with DALI, we recovered weak but significant (Z=2.6) hits from long LmuC to the C-terminal region of SbcD of *E. coli*. This partial match was further confirmed by using FoldSeek with different homologs of long LmuC (Best hit probability P=1, **Extended Data Fig. S16).** This striking observation suggests that long LmuC derived from SbcD. For LmuA short and long CTDs, we could not find similarity to other known domains. In contrast, many effectors like Cap4, Mrr, Protease and Sir2 were also found in other defense systems, especially in the Avs family. Contrarily to LmuC, we could not find evidence of homology between LmuA and SbcCD.

Since long LmuB are close homologs of SbcC, and long LmuC are structurally similar to the SbcC-binding domain of SbcD (SbcD CTD), we hypothesized that Lamassu evolved from an ancient SbcCD system. The operon architecture of Lamassu is compatible with such a scenario, since the SbcD CTD lies just before SbcC, like LmuC before LmuB (**Fig. 5d**). We thus studied the conservation of the structure and molecular complex using structural model predictions. We generated 20 complexes *in silico*, including the SbcCD system of *E. coli*, long (n=5) and short Lamassus (n=14) (**Extended Data Fig. S17**). Complexes of long and short were more different between long and short than within, revealing only two major conserved architectures for both. Short LmuC were positioned similarly as in the Lamassu Vc-Cap4 complex, while long LmuC had an additional short domain separated by an unstructured linker (**Fig. 5c**). Similarly, short LmuA CTDs are predicted to form a conserved bundle of 5-6 helices, while long LmuA CTDs are predicted to form a more complex fold with helices, a conserved loop with negatively charged residues (predicted to bind LmuB), and a beta-sheet (**Fig. 5c**). In predicted complexes of long Lamassu, LmuA and LmuC were predicted to interact around the head domains of LmuB, reminiscent of the interaction between SbcC and SbcD ^29^. Strikingly, long LmuC were positioned similarly to the CTD of SbcD.

We propose an evolutionary model **(Fig. 5a,b**) for short Lamassu systems originating from an ancestral SbcCD complex with long coiled-coil domains, standard Walker motifs, and a single CxxC motif. Separation of SbcD’s nuclease and CTD gave rise to LmuC and facilitated modular effector integration. Long Lamassu evolved through inovation of the LmuA CTD, retaining key motifs but shortening coiled-coil regions by ∼200 amino acids. Short Lamassu further compacted from long Lamassu by additional coiled-coil reduction and significant divergence or innovation of the LmuA CTD and LmuC. These short variants display diversified Walker B motifs, greater effector diversity, and in some cases, loss of entire components (LmuC or effectors). This staged reduction underscores the gradual remodeling of a DNA-repair system into a defense complex.

## DISCUSSION

Our study reveals how Lamassu evolved from a DNA repair ancestor through major structural innovations that enabled its transition into an antiphage defense system. Based on our results, we propose that as the LmuABC complex scans along dsDNA, it recognizes and binds free DNA ends or a related DNA substrate, with this binding event mediated by LmuB and LmuC. We postulate that ATP hydrolysis could promote the disruption of the LmuABC-DNA complex, facilitating LmuA release and tetramerization into an activated state. This active tetramer mediates widespread, non-specific cleavage of cellular dsDNA. The need for disengagement of LmuA from the LmuABC complex for activation through changes in the oligomerization status could be a conserved mechanism across different Lamassu systems as several LmuA effectors are predicted to work in an oligomeric state.

Lamassu systems fall into two families—long and short forms—both built around a conserved SMC-like LmuB scaffold. While LmuA-NTDs are often homologous, the LmuA-CTDs and LmuC proteins differ markedly, reflecting dynamic modular evolution likely enabling adaptation to diverse phage threats. By solving the apo and dsDNA-bound structures of the short-form Lamassu Vc-Cap4, we reveal that it forms an atypical SMC-like complex, distinct from both canonical SMC proteins and RAD50/SbcC-like complexes. Unlike the expected rod-shaped dimer with partner proteins interacting around the head domains, LmuB adopts a kinked coiled-coil conformation, with its apex contacting LmuC and the CTD of LmuA. Both LmuB heads are not visible in the apo-LmuABC structure suggesting flexibility, with one of them (LmuB1) visible on binding to dsDNA. We suggest recognition of DNA ends leads to disengagement of the LmuB2 head, while only the LmuC-bound head interacts with DNA. This disengaged state correlates with activation, likely driven by ATP hydrolysis, as ATP**γ**S does not stimulate activation and the LmuB K40A mutant is inactive. Functional assays confirm that interactions between LmuB, LmuA and LmuC are essential for activity in vivo.

The Lamassu LmuABC complex recognizes double-stranded DNA ends in vitro, a feature explained by structural elements in LmuB that block duplex from threading through the complex. These results are in line with a recent preprint on a close homolog of Lamassu Vc-Cap4 ^30^. LmuC also interacts with the dsDNA, suggesting it may function as a specificity factor that stabilizes the interaction between LmuB and DNA ends or a structurally related substrate. In uninfected cells, we suggest LmuB could continuously scan genomic DNA in search of its cognate substrate. Upon infection with Lambda-vir, a strong and specific enrichment of LmuB at the phage’s origin of replication is observed. This mode of replication continues until approximately 16 minutes post-infection ^31^, which aligns with our observation that LmuB is enriched at the origin at 10 minutes, but not at 20 minutes post-infection—when replication has transitioned to the rolling-circle mode ^31^.

We propose that Lamassu senses DNA structures specific to phage replication, which could derive from iterons. Supporting this, Lamassu is triggered by structured DNA motifs like palindromes and can be blocked by host or phage mutations affecting replication initiation^11^. We observe that LmuB specifically binds iterons in the O gene in a replication context. Those iterons are predicted to form hairpin or cruciform structures^32^, and are bound to the O protein. Formation of the O-iterons complex leads to DNA bending that facilitates the unwinding of an adjacent AT-rich sequence ^33–35^ allowing bidirectional theta replication to proceed ^22,33,34^. These findings suggest that Lamassu detects unstable DNA intermediates—such as open origins or stalled forks—associated with phage replication, allowing it to distinguish foreign from host DNA and act as a sensor of “reckless” replication events. In this way, Lamassu may have evolved not simply to recognize replication per se, but to discriminate the timing, coordination, and molecular signatures that distinguish phage replication from its host counterpart.

Beyond DNA recognition, our study reveals a major conformational transition in LmuA, linked to activation. In the LmuA₁B₂C₁ complex, LmuA is monomeric and extended, stabilized by interactions with LmuB and LmuC. However, when excess LmuA is added to LmuBC, forming an activated complex in biochemical assays, LmuA assembles into a tetrameric state observed by cryo-EM, in line with recent observations n a related system ^30^. This tetramer stabilizes two Cap4 NTDs, a distant relative of the PD-(D/E)xK nuclease family also associated to CBASS and Avs systems ^36,37^, into a single active cleavage pocket, while the remaining two Cap4 NTDs do not engage in cleavage. ATP hydrolysis most likely provides the energy to trigger the conformational changes necessary for disengagement of LmuA from LmuABC and thus LmuA activation by oligomerization via its CTD.

The evolutionary trajectory of the Lamassu system suggests it originated from the DNA repair complex SbcCD, with long Lamassu being an intermediate between this ancestral repair machinery and the compacted short Lamassu variants. This transition likely involved repurposing SbcCD’s “detect-and-repair” function into a “detect-and-kill” strategy, shifting from site-specific degradation of DNA to an abortive infection system relying on promiscuous effectors. Unlike SbcCD, which precisely positions SbcD for cleavage, Lamassu activates diverse effectors, including nucleases like LmuA Cap4 and enzymes such as Sir2, proteases, and FMO, broadening its defensive potential. Beyond structural and sequence similarities between SbcCD and Lamassu is their shared substrate recognition: SbcCD detects and processes palindromic DNA, excising these sequences and resolving replication fork structures ^25^, while the human orthologs Rad50-Mre11 detects dsDNA ends ^26^. The ability to recognize structural DNA motifs rather than specific sequences suggests that Lamassu evolved by co-opting an ancestral DNA surveillance function, transforming it from a genome maintenance tool into a broad-spectrum immune defense system.

Many proteins involved in antiphage defense systems contain domains that are also found in DNA repair proteins, highlighting a deep evolutionary link between genome maintenance and immunity ^38^. For instance, the RecB-like nuclease of RecBCD is co-opted in Cas4 for CRISPR spacer acquisition^39^, and RecBCD itself degrades foreign DNA ^40^, countered by phage-encoded inhibitors like Lambda Gam. Other repair domains—Toprim, Ku, and helicase motifs from proteins like RecQ—also appear in defense systems^41;42,9^, highlighting a recurrent evolutionary link between genome maintenance and immunity. This evolutionary theme extends beyond bacteria, as eukaryotic cells also exploit DNA repair complexes for antiviral defense. For example, RAD50, part of the MRN complex, partners with CARD9 to sense cytoplasmic viral DNA^43^ while the Smc5/6 complex restricts viruses but is targeted for degradation by Hepatitis B virus. ^44,45^. These examples illustrate that DNA repair enzymes have repeatedly served as a reservoir for the evolution of immune defenses, reflecting a broader evolutionary strategy where existing molecular machineries are repurposed for defense against pathogens.

## MATERIALS AND METHODS

### Genome database

We used the RefSeq complete genomes database from the National Center for Biotechnology Information (NCBI), downloaded in July 2022, composed of 22,920 prokaryotic genomes. Genomes were formatted in the gembase format with PaNaCota v1.3 ^46^ and annotated with DefenseFinder v1.09 ^47–49^ and DefenseFinder models v1.1.0 ^47–49^. The list of genomes is available in **Extended Data Table 2**.

### Lamassu detection

To comprehensively identify Lamassu systems, we first assembled a dataset incorporating protein sequences from experimentally validated Lamassu systems from Doron *et al.* ^7^, Millman *et al.* ^9^, the ddmABC system ^10,11,20^, as well as predicted systems from Krishnan *et al.* ^8^(ABC-3C). LmuB is the only gene that can be aligned between all previously identified systems, so we used this alignment to build a seed profile HMM of LmuB. We used this seed profile to search across 22,920 complete genomes (See *Genome database*). All significant hits (E-value < 1e-3) were recovered (n=367,581) and clustered at 30% identity and coverage with MMseqs2 v14.7e284 ^50^. After removing singleton families, we aligned the representative sequences of the 895 remaining families and built a phylogeny. This yielded a star-like phylogenetic tree with deep branching clades corresponding to both known and unknown genes based on gene annotation (**Extended Data Fig. S15**). Known genes comprised SbcC homologs, ABC-transporters, and some defense systems included in DefenseFinder like DndD or RloC. By analyzing the genetic neighborhood of the representative sequences, we identified two deep branching clades often associated with previously identified LmuC and LmuA homologs and hypothesized that those two clades represented *bona fide* Lamassus systems (**Extended Data Fig. S15**). We thus selected both clades, retrieved the cluster members (n=3786) and analyzed their genomic context.

We retrieved 75,425 genes surrounding the putative LmuB identified in the previous step and extracted all operons of two to five genes based on the distance between genes (<30bp apart) and direction (same direction). This retrieved 5,892 genes, which were annotated with Pfam and DefenseFinder profile HMMs, and clustered into 379 families. Operons were automatically included in our set of putative Lamassu if they included at least one domain previously associated with Lamassu (3C-ABC domains, Lamassu DefenseFinder genes). For the remaining families, we filtered out the singleton families and manually curated the remaining families by re-annotating then with HH-pred Pfam which detects more remote homologs. This approach identified 2646 Lamassu systems which were later used to build individual profile HMM (see *DefenseFinder models*). Defense scores were computed using a procedure adapted from Doron *et al* ^7^. For each 90% coverage and identity protein families, we compute the fraction with at least one defense system (as detected by DefenseFinder) in its genomic neighborhood of +/- 10 genes around. Then we average the score for all the members of the gene family to obtain the defense-score of the family.

### Clustering and alignments

Set of protein sequences were routinely analyzed with the following method, unless stated otherwise: sequences were first clustered with Mmseqs2 at 80% sequence identity and 80% bidirectional coverage, and clusters with less than 3 members were excluded as they mainly contained truncated protein sequences. Representative sequences of each cluster were aligned with Mafft v7.526 ^51^(GINSI algorithm, unless otherwise stated). Alignments were manually inspected for correct alignment of known motifs of catalytic domains and structural features with Unipro UGENE v51^52^ and mis-aligned and heavily truncated sequences were removed before re-producing a novel alignment without including them. FoldMason ^53^ was used to generate multiple sequence alignments from AF3 models. This is especially useful to align structurally conserved regions with high degree of sequence divergence, like the coiled-coils region of LmuB.

### Phylogeny

Phylogenetic trees were computed with Iq-tree v2.2^54^ and ModelFinder ^55^ with the following options -mset WAG,Q.pfam -B 1000. Trees were visualized and annotated with TreeViewer v2.2.0 ^56^. When an alignment was too large for efficient phylogenetic reconstruction, we used Clipkit v2.3 ^57^ to filter out columns with >90% gap. This is indicated by the _g90 suffix in the alignment and tree files.

### DefenseFinder models

We retrieved the sequences of the 2646 systems identified by our detection method (see Lamassu Detection) and assigned them a gene name based on their annotation, length and position in the operons. We then clustered LmuA and LmuC proteins, aligned their representative sequence and we attempted to build phylogenies. This yielded star-like phylogenies with deep-branching clades, indicative that not all proteins could be aligned properly. We thus selected deep-branching clades (n=18 for LmuA, n=12 for LmuC), re-aligned them independently and built profile HMM with Hmmer.

We then built a set of models tailored to detect Lamassu type I and type II, short and long, with the effector. While giving priority to our novel profiles, we also added the ones built in the previous iteration of DefenseFinder and by Krishnan et al ^8^. We set up tailored gathering thresholds (Cut-ga) for each profile HMM, based on the distribution of hits in close proximity to the other components, as previously described ^47^. Our models can detect the sub-systems presented in this work, but may also find new ones which were not present in the databases we used. This is possible because we created a generalist model with LmuB and LmuC profiles, as well as LmuA CTD profiles of Krishnan et al ^8^. This approach enabled us to retrieve n=3,829 Lamassu systems. These new models and definitions are available at https://github.com/mdmparis/defense-finder-models. The list of detected systems is available as **Extended Data Table S3**.

### Strains and plasmids

Strains and plasmids used in this study are available as **Extended Data Table S4**.

### Phage plaque assays

Phage plaque assays were conducted using LB-Lennox agar supplemented with 0.7% agar, 100 μM carbenicillin (Carb), 5 mM MgSO₄, and 5 mM CaCl₂. Varying concentrations of L-arabinose were included to induce the expression of the systems. To prepare the bacterial lawn, 300 μL of overnight bacterial cultures grown in LB-Miller medium supplemented with 100 μM carbenicillin and 0.02% glucose were mixed with 20 mL of LB-Lennox agar. The mixture was poured onto plates and allowed to solidify for 30 minutes at room temperature.

Phage spotting was performed by applying 5 μL of serially diluted phage suspensions in LB-Lennox medium onto the solidified bacterial lawn. Plates were incubated at 37°C for 16–18 hours before imaging the resulting plaques. The last phage dilution showing lysis on the plate was reported and compared with the associated negative control (pBAD-RFP). Efficiency of plating is the ratio of last dilution showing lysis of the experiment over the control. An efficiency of plating of 1 means there is no protection.

### Mutagenesis

Mutagenesis was performed either by Genscript or using the Q5 Site-Directed Mutagenesis Kit (New England Biolabs, E0554S) according to the manufacturer’s protocol. The reactions were optimized to ensure efficient amplification. Mutagenic primers were designed with NEBaseChanger webserver, ensuring optimal annealing temperatures and minimal secondary structures. All primers used for mutagenesis are listed in Extended Data Table S3.

Briefly, Following PCR amplification, the reactions were treated with kinase, ligase, and DpnI (KLD) enzyme mix to circularize the mutated plasmids and degrade the parental template DNA. The resulting products were transformed into chemically competent Escherichia coli DH10 Beta cells, and transformants were selected on LB-agar plates containing the appropriate antibiotic. To confirm the mutations, minipreps were sequenced by whole-plasmid sequencing using PlasmidSaurus long-read sequencing. Sequencing results were analyzed to verify the presence of the desired mutations and to ensure no off-target changes were introduced. Primers sequences **Extended Data Table S5**.

### Structural modeling

We used AlphaFold3^58^ with a fixed stoichiometry deduced from the nMs data and cryo-EM structures for comparative structural studies. 1 LmuA, 1 LmuC, 2 LmuB with 2 Mg2+, 1 Zn2+, 2 ADP and 72-bp dsDNA. The best ranking models were evaluated and visualized with ChimeraX v1.9 ^59^. We used the match command to compute the root-mean-square deviation (RMSD) between models. We used the matchmaker function with the secondary structure weighting set to 1 to superimpose distant domains such as LmuA CTD and whole-proteins such as LmuB and LmuC. We used the DALI webserver ^60^ to compute pairwise similarity as a Z-score between all models.

### Chromatin immunoprecipitation sequencing (ChIP-Seq)

For all ChIP experiments, an active site mutation (K57A) was introduced into the LmuA gene to create pLmuABC to prevent Cap4-mediated degradation. LmuB was 3×Flag-tagged while the other proteins remained untagged. Transformations of the system and if applicable, the plasmid encoded binding site pTarget, were performed according to the general transposition assay protocol. Single colonies were picked and grown overnight with 2% glucose and antibiotics. Cultures were then spun down and washed to remove glucose by resuspension in liquid LB before being diluted 1/100 in 26 mL fresh media with antibiotics. At OD600 = .1, cultures were induced with .02% arabinose. For phage infected samples, cultures were grown to OD600 = .9, before lambda phage infection at multiplicity of infection (MOI) 5. Samples were collected at 5, 10, and 20 minute time points post infection. Plasmid based samples were grown to OD600 = .6 before collection.

Cross-linking and immunoprecipitation were generally performed according to previously established protocols ^61^. All samples were first normalized to 40 mL culture volume before adding formaldehyde (1 ml of 37% solution; Thermo Fisher Scientific) and mixed immediately by vortexing. Cross-linking was performed by gently shaking at room temperature for 20 min. To stop cross-linking, 4.6 ml of 2.5 M glycine (∼0.25 M final concentration) were added, followed by 10 min incubation with gentle shaking. Cells were pelleted at 4 °C by centrifuging at 4,000g for 8 min. The following steps were performed on ice using buffers that had been sterile-filtered through a 0.22 μm filter. The supernatant was discarded and the pellets were fully resuspended in 40 ml TBS buffer (20 mM Tris-HCl pH 7.5, 0.15 M NaCl) by vortexing. After centrifuging again at 4,000g for 8 min at 4 °C, the supernatant was removed, and the pellet was again resuspended in 40 ml TBS buffer. Next, the OD600 was measured for a 1:1 mixture of the cell suspension and fresh TBS buffer, and a standardized volume equivalent to 40 ml of OD600 = 0.6 was aliquoted into new 50 ml conical tubes. A final 8 min centrifugation step at 4,000g and 4 °C was performed, cells were pelleted and the supernatant was discarded. Residual liquid was removed by briefly inverting the tube, and cell pellets were flash-frozen using liquid nitrogen and stored at −80 °C or kept on ice for the subsequent steps.

Bovine serum albumin (GoldBio) was dissolved in 1× PBS buffer (Gibco) and sterile-filtered to generate a 5 mg ml−1 BSA solution. For each sample, 25 μl of Dynabeads Protein G (Thermo Fisher Scientific) slurry (hereafter, beads or magnetic beads) were prepared for immunoprecipitation. Beads from up to 250 μl of the initial slurry were processed together in a single tube, and washes were performed at room temperature, as follows: the slurry was transferred to a 1.5 ml tube and placed onto a magnetic rack until the beads had fully settled. The supernatant was removed carefully, 1 ml BSA solution was added, and the beads were fully resuspended by vortexing, followed by rotating for 30 s. This was repeated for three more washes. Finally, the beads were resuspended in 25 μl (× n samples) of BSA solution, followed by addition of 4 μl (× n samples) of monoclonal anti-Flag M2 antibodies produced in mouse (Sigma-Aldrich). The suspension was moved to 4 °C and rotated for >3 h to conjugate antibodies to the magnetic beads. While conjugation was proceeding, cross-linked cell pellets were thawed on ice, resuspended in FA lysis buffer 150 (50 mM HEPES-KOH pH 7.5, 0.1% (w/v) sodium deoxycholate, 0.1% (w/v) SDS, 1 mM EDTA, 1% (v/v) Triton X-100, 150 mM NaCl) with protease inhibitor cocktail (Sigma-Aldrich) and transferred to a 1 ml milliTUBE AFA Fiber (Covaris). The samples were sonicated on a M220 Focused-ultrasonicator (Covaris) with the following SonoLab 7.2 settings: minimum temperature, 4 °C; set point, 6 °C; maximum temperature, 8 °C; peak power, 75.0; duty factor, 10; cycles/bursts, 200; 17.5 min sonication time. After sonication, samples were cleared of cell debris by centrifugation at 20,000g and 4 °C for 20 min. The pellet was discarded, and the supernatant (∼1 ml) was transferred into a fresh tube and kept on ice for immunoprecipitation. For non-immunoprecipitated input control samples, 10 μl (∼1%) of the sheared cleared lysate were transferred into a separate 1.5 ml tube, flash-frozen in liquid nitrogen and stored at −80 °C.

After >3 h, the conjugation mixture of magnetic beads and antibodies was washed four times as described above, but at 4 °C. Next, the beads were resuspended in 30 μl (× n samples) FA lysis buffer 150 with protease inhibitor, and 31 μl of resuspended antibody-conjugated beads were mixed with each sample of sheared cell lysate. The samples were rotated overnight for 12–16 h at 4 °C for immunoprecipitation of Flag-tagged proteins. The next day, tubes containing beads were placed on a magnetic rack, and the supernatant was discarded. Then, six bead washes were performed at room temperature, as follows, using 1 ml of each buffer followed by sample rotation for 1.5 min: (1) two washes with FA lysis buffer 150 (without protease inhibitor); (2) one wash with FA lysis buffer 500 (50 mM HEPES-KOH pH 7.5, 0.1% (w/v) sodium deoxycholate, 0.1% (w/v) SDS, 1 mM EDTA, 1% (v/v) Triton X-100, 500 mM NaCl); (3) one wash with ChIP wash buffer (10 mM Tris-HCl pH 8.0, 250 mM LiCl, 0.5% (w/v) sodium deoxycholate, 0.1% (w/v) SDS, 1 mM EDTA, 1% (v/v) Triton X-100, 500 mM NaCl); and (4) two washes with TE buffer 10/1 (10 mM Tris-HCl pH 8.0, 1 mM EDTA).

The beads were then placed onto a magnetic rack, the supernatant was removed and the beads were resuspended in 200 μl of ChIP elution buffer (1% (w/v) SDS, 0.1 M NaHCO3) made fresh on the day of the washes. To release protein–DNA complexes from beads, the suspensions were incubated at 65 °C for 1.25 h with gentle vortexing every 15 min to resuspend settled beads. During this incubation, the non-immunoprecipitated input samples were thawed, and 190 μl of ChIP Elution Buffer was added, followed by the addition of 10 μl of 5 M NaCl. After the 1.25 h incubation of the immunoprecipitated samples was complete, the tubes were placed back onto a magnetic rack, and the supernatant containing eluted protein–DNA complexes was transferred to a new tube. Then, 9.75 μl of 5 M NaCl was added to ∼195 μl of eluate, and the samples (both immunoprecipitated and non-immunoprecipitated controls) were incubated at 65 °C overnight (without shaking) to reverse-cross-link proteins and DNA. The next day, samples were mixed with 1 μl of 10 mg ml−1 RNase A (Thermo Fisher Scientific) and incubated for 1 h at 37 °C, followed by addition of 2.8 μl of 20 mg ml−1 proteinase K (Thermo Fisher Scientific) and 1 h incubation at 55 °C. After adding 1 ml of buffer PB (QIAGEN recipe), the samples were purified using QIAquick spin columns (QIAGEN) and eluted in 40 μl TE buffer 10/0.1 (10 mM Tris-HCl pH 8.0, 0.1 mM EDTA).

Illumina libraries were generated for immunoprecipitated and input samples using the NEBNext Ultra II DNA Library Prep Kit for Illumina (NEB). Sample concentrations for ChIP–seq were determined using the DeNovix dsDNA Ultra High Sensitivity Kit. Starting DNA amounts were standardized such that an approximately equal mass of all input and immunoprecipitated DNA was used for library preparation. After adapter ligation, a single PCR amplification was performed to add Illumina barcodes, and ∼450 bp DNA fragments were selected using two-sided AMPure XP bead (Beckman Coulter) size selection, as follows: the volume of barcoded immunoprecipitated and input DNA was brought up to 50 μl with TE Buffer 10/0.1; in the first size-selection step, 0.55× AMPure beads (27.5 μl) were added to the DNA, the sample was placed onto a magnetic rack, and the supernatant was discarded and the AMPure beads were retained; in the second size-selection step, 0.35× AMPure beads (17.5 μl) were added to the DNA, the sample was placed onto a magnetic rack, and the AMPure beads were discarded and the supernatant was retained. The concentration of DNA was determined for pooling using the DeNovix dsDNA High Sensitivity Kit.

Illumina libraries were sequenced in either single- or paired-end mode on the Illumina MiniSeq and NextSeq platforms with automated demultiplexing and adapter trimming (Illumina). For each ChIP– seq sample, >2,000,000 raw reads (including genomic and plasmid-mapping reads) were obtained.

### LmuBC protein expression and purification

The genes encoding LmuB (UniProt ID: A0A060KSR0) and LmuC (UniProt ID:B7X6T6) from *V. cholerae* serotype O1 biovar El Tor were codon optimized and synthesized by IDT. They were then cloned into a pRSF-Duet vector with Kanamycin resistance with LmuB in multiple cloning site 1 and LmuC in multiple cloning site 2. LmuB was fused to a His-SUMO tag, while LmuC was expressed without a tag.

N-His SUMO LmuB and LmuC were then co-expressed in BL21(DE3) cells with induction by 0.4 mM IPTG (GoldBio) at OD 0.4-0.6, followed by shaking at 16 °C for 16 hours. The cells were harvested by centrifugation at 22.000 rpm for 1 hr followed by resuspension in lysis buffer (50 mM Tris pH 8, 500 mM NaCl, 5% glycerol, 5 mM b-mercaptoethanol, complete EDTA-free protease inhibitor, benzonase). Resuspended cells were lysed through sonication (1s pulse on and 2s pulse off time) and centrifuged at 113092g at 4 °C for 1 hour. Supernatant was then passed through a prepacked Ni NTA column (Cytiva Life Sciences™ HisTrap™ FF). This was followed by washing the column with Wash buffer 1 (50 mM Tris pH 8, 1 M NaCl, 5% glycerol, 5 mM b-mercaptoethanol) and then by the lysis buffer. Protein was then eluted using 50 mM Tris pH 8, 500 mM NaCl, 5% glycerol, 5 mM b-mercaptoethanol and 500 mM Imidazole using gradient elution using AKTA-FPLC (AKTA pure^TM^ chromatography system) to separate out impurities. Pure fractions were then subjected to an overnight Ulp1 protease digestion at 4 °C. These fractions were then pooled, and salt reduced to 250 mM by addition of 50 mM Tris pH 8, 20 mM NaCl, 5% glycerol and 2 mM b-mercaptoethanol (buffer A) before loading onto a Heparin column (HiTrap Heparin HP, cytiva) to remove contaminating DNA. Protein was eluted by using a gradient using buffer A (50 mM Tris pH 8, 20 mM NaCl, 5% glycerol and 2 mM b-mercaptoethanol) and buffer B (50 mM Tris pH 8, 1 M NaCl, 5% glycerol and 2 mM b-mercaptoethanol) with the AKTA-FPLC system. Pure fractions were then combined and concentrated with centricon-plus 70 centrifugal filter 30 kDa (Millipore) for loading onto a superdex 200 Increase 10/300 GL column (GE Healthcare) equilibrated with SEC buffer (25 mM Hepes, pH 7.5, 250 mM NaCl and 5 mM DTT). Peak fractions were then collected, concentrated and prepared for SDS-PAGE and cryo-EM analysis.

### LmuA expression and purification

The genes encoding LmuA (UniProt ID: A0A060KT36) from *V. cholerae* serotype O1 biovar El Tor was codon optimized and synthesized by IDT. It was then cloned into a pRSF-Duet vector with Kanamycin resistance and fused to a His-SUMO tag. The gene was then expressed in BL21-DE3 followed by induction by 0.4 mM IPTG (GoldBio) at OD 0.4-0.6, followed by shaking at 16°C for 16 hours. The cells were harvested by centrifugation followed by resuspension in lysis buffer (50mM Tris pH 8, 500 mM NaCl, 5% glycerol, 5mM β-mercaptoethanol, complete EDTA-free protease inhibitor, benzonase). Resuspended cells were lysed through sonication and centrifuged at 113092 g at 4°C for 1 hour. Supernatant was then passed through a prepacked Ni NTA column (Cytiva Life Sciences^TM^ HisTrap^TM^ FF). This was followed by washing the column with wash buffer 1 (30mM Imidazole, 50mM Tris pH 8, 1M NaCl, 5% glycerol, 5 mMβ-mercaptoethanol) and then by lysis buffer. Protein was then eluted using gradient elution using AKTA-FPLC (AKTA pure^TM^ chromatography system) to separate out impurities. Pure fractions were then subjected to an overnight Ulp1 protease digestion at 4°C. These fractions were then pooled, and the salt reduce to 250 mM before loading onto a Heparin column (HiTrap Heparin HP, cytiva) to remove contaminating DNA. Protein was eluted by a gradient using buffer B (50 mM Tris pH 8, 1M NaCl, 5% glycerol and 2 mM β-mercaptoethanol) with the AKTA-FPLC system. Pure fractions were then combined and concentrated for loading onto a superdex 200 Increase 10/300 GL column (GE Healthcare) equilibrated with SEC buffer (25 mM Hepes, pH 7.5, 250 mM NaCl and 5 mM DTT). Peak fractions were then collected, concentrated and prepared for SDS-PAGE and cryo-EM analysis. To generate K57A mutation, primers were designed and purchased following subsequent cloning using PCR. Both the wild type and mutant LmuA were overexpressed in BL-21 (DE3) cells and purified using the same set of conditions as LmuBC except 25 mM Tris, pH 8, 500 mM NaCl and 5 mM DTT was employed to perform size-exclusion chromatography. Notably, we observed severe aggregation of both LmuA and LmuA (K57A) on the sizing column (Superdex. 200 Increase 10/300 gl). Therefore, we directly used LmuA and Lmu(K57A) from the heparin column for LmuABC complex generation.

### Sample and cryo-EM grid preparation, data collection, processing and model building for the LmuA tetramer

We prepared a 0.3 mg/ml sample containing LmuA (from the heparin column) and LmuBC (from a sizing column) at a 5:1 ratio of LmuA: LmuBC prior to grid freezing without any cross-linking agent. A 4 µl aliquot of the sample was applied to graphene oxide grids (R1.2/1.3 AU 400 Mesh). The grids were blotted for 1 second with a force setting of 0 at 4 °C and 100% humidity, then rapidly frozen in liquid ethane using a Vitrobot Mark IV (FEI).

Data collection was performed at the Memorial Sloan Kettering Cancer Center (MSKCC) using a Titan Krios G2 transmission electron microscope (FEI) operating at 300 kV and equipped with a K3 direct detector, controlled by SerialEM software. Movies were recorded in super-resolution mode with a total electron dose of 53 e^−^/Å², a defocus range of -0.8 to -2.2 µm, and a pixel size of 1.064 Å. For structure determination, a total of 7,002 movies was processed using cryoSPARC, Patch motion correction and patch contrast transfer function (CTF) estimation were applied to correct for drift and estimate CTF parameters, respectively. Micrographs with ice contamination, high astigmatism, or poor CTF fit resolution were excluded by setting appropriate threshold ranges using the ’Manually Curate Exposures’ job. Particles from high-quality micrographs were then picked with the Blob picker and extract micrograph jobs, resulting in the selection of 5,768,112 particles. Multiple rounds of two-dimensional (2D) classification were conducted to eliminate junk particles. After several rounds of ab-initio reconstruction and heterogeneous refinement followed by further refining through homogeneous and non-uniform refinement yielded two conformations of an LmuA tetramer. Model building was guided by a LmuA tetramer fold predicted using AlphaFold3 ^58^, followed by structure building in Coot ^62^, with the atomic coordinates refined against the map in PHENIX ^63^. These efforts yielded a pair of LmuA tetramer structures only, despite the presence of minor proportion of LmuBC in sample preparation. The two-fold symmetric LmuA tetramer based on 193,953 particles was solved at a resolution of 3.13 Å, while the LmuA tetramer lacking symmetry based on 197,714 particle was solved at a resolution of 3.4 Å, which enabled precise model building for both structures. In order to filter the LmuA symmetrical and asymmetrical tetramer maps, we used the sharp.mrc file in Chimera. We next applied the ‘Volume Filter’ with a width of 1.2 (approximately matching the pixel size) and selected the Gaussian filter type. The filtered maps were saved for subsequent analysis and Figure preparations. The resolution estimates were based on a Fourier shell correlation (FSC) cutoff of 0.143. Figures were prepared using UCSF Chimera ^64^ and USCF ChimeraX ^59^ with the final image layout created in Adobe Illustrator.

### Sample and cryo-EM grid preparation, data collection, processing and model building for apo-LmuABC

Following separate LmuBC and LmuA purifications from their respective heparin columns, both were mixed at 1:5 ratio of LmuBC:LmuA and incubated overnight at 4 °C. These fractions were then concentrated and passed through a Superdex 200 Increase 10/300 GL column (GE Healthcare) to generate apo-LmuABC using 25 mM Hepes, pH 7.5, 250 mM NaCl and 5 mM DTT as the equilibration and elution buffer. The protein was then concentrated to 0.6 mg/ml, followed by 0.025% glutaraldehyde incubation on ice for 30 minutes for cross-linking. Following spinning down to remove precipitates, the reaction was stopped by the addition of 0.1 M Tris, pH 8. A 4 µl aliquot of apo-LmuABC in the presence 1 mM PPi was applied to glow-discharged holey gold grids (UltrAuFoil 300 mesh R1.2/1.3) with a blotting time of 2 s and wait time of 10 s.

Data collection for apo-LmuABC was performed at MSKCC on a Titan Krios G4 transmission electron microscope (FEI) operating at 300 kV and equipped with a Falcon 4i direct detector, controlled by EPU software. Movies were recorded in counting mode with a total electron dose of 29.28 e^−^/Å², a defocus range of -0.8 to -2.2 µm, and a pixel size of 0.725 Å. For LmuABC structure determination, 18,780 movies were processed using the same procedure outlined above. Following particle selection through Blob picking and template-based methods, 1,683,512 particles were chosen for 2D classification. After several rounds of 2D classifications to ensure best particles were selected, several rounds of ab-initio and heterogenous refinements were performed to generate initial models and enhance density map quality. The best class from the heterogeneous refinement with 127,014 particles was selected and further polished using homogeneous and non-uniform refinement. Model building was guided by LmuA, LmuB and LmuC folds predicted using AlphaFold3 ^58^, followed by structure building in Coot ^62^, with the atomic coordinates refined against the map in PHENIX ^63^. This process resulted in a structure of apo-LmuA_1_B_2_C_1_ at 3.28 Å resolution, allowing for accurate model building. The resolution estimation was based on a Fourier shell correlation (FSC) cutoff of 0.143.

### Sample and cryo-EM grid preparation, data collection, processing and model building for Lmu(K57A)ABC-DNA complex

A sample of LmuA(K57A)BC was prepared using 1:2 ratio of LmuBC:LmuA(K57A) from their respective heparin columns, incubated overnight at 4 °C and then run on size-exclusion column to get a peak fraction, to which was added 20-bp dsDNA (GTGATAGTTAGAAACGTAAT sequence and its complementary strand). The complex was concentrated to 0.6 mg/ml at a protein:DNA ratio of 1:4 and incubated for 1 hour on ice, followed by cross-linking with 0.025% glutaraldehyde on ice for 30 minutes. A 4 µl aliquot of LmuA(K57A)BC-DNA complex in the presence 1 mM PPi was applied to glow-discharged holey gold grids (UltrAuFoil 300 mesh R1.2/1.3) with a blotting time of 2 s and wait time of 10 s.

Data collection for LmuA(K57A)BC-DNA complex was performed at MSKCC on a Titan Krios G4 transmission electron microscope (FEI) operating at 300 kV and equipped with a Falcon 4i direct detector, controlled by EPU software. Movies were recorded in counting mode with a total electron dose of 29.28 e–/Å², a defocus range of -0.8 to -2.2 µm, and a pixel size of 0.725 Å. For structure determination of the LmuA(K57A)BC-DNA complex, a total of 11,653 movies were processed using cryoSPARC, Patch motion correction and patch contrast transfer function (CTF) estimation were applied to correct for drift and estimate CTF parameters, respectively. Micrographs with ice contamination, high astigmatism, or poor CTF fit resolution were excluded by setting appropriate threshold ranges using the ’Manually Curate Exposures’ job. High-quality micrographs were then processed with the Blob picker and extract jobs, resulting in the selection of 2,384,679 particles. Multiple rounds of two-dimensional (2D) classification were performed to remove junk particles. Following two rounds of ab initio reconstruction and heterogeneous refinement into three classes, with two of these corresponding to monomeric apo-LmuA(K57A)BC and monomeric LmuA(K57A)BC-dsDNA complex. These volumes were further refined using homogeneous and non-uniform refinement, followed by structure building in Coot ^62^, with the atomic coordinates refined against the map in PHENIX ^63^. For the monomeric apo-LmuA(K57A)1B2C1 reconstruction, 135,737 particles were used, achieving a resolution of 3.21 Å, enabling precise model building. For the monomeric LmuA(K57A)_1_B_2_C_1_-DNA complex, 190,887 particles were utilized, reaching a resolution of 2.93 Å. The third class reflecting formation of a dimeric LmuA(K57A)_1_B_2_C_1_-DNA complex contained only 2,690 particles and hence the structure could only be determined to a resolution of 11.2 Å. All resolution estimations were based on a Fourier shell correlation (FSC) cutoff of 0.143.

### *In vitro* DNA degradation experiments

For the *in vitro* ATP dependent DNA degradation experiments (**Fig. 4a)**, 40 nM pUC19 vector (Thermo Scientific) was incubated with 20 nM LmuABC. The reaction was conducted in the reaction buffer (RB) containing 50 mM Hepes pH 7.5, 50 mM KCl, 1 mM DTT, 10 mM MgCl_2_ (RB) with increasing concentrations of ATP ranging from 0.5 to 3 mM. ATP**γ**S was added to the final reaction at 3 mM concentration to check if ATP binding and subsequent hydrolysis activated LmuABC-catalyzed DNA cleavage. All assays were conducted at 37 °C for 1 hour and stopped by addition of 1 mg/ml of proteinase K (Thermo Scientific) incubated at 55°C for 10 minutes followed by loading it in 1% agarose gel (stained with SYBR Safe, Thermo Fischer) with 1X purple loading dye (New England Biolabs).

For the *in vitro* ATP dependent DNA degradation experiments in the presence of 20-bp dsDNA (**Fig. 4b)**, the reactions were performed similar to ATP dependent reactions with the same reaction buffer but with LmuABC preincubated with 0.2μM20-bp dsDNA.

For the concentration dependency experiment (**Extended Data Fig. 12a)**, 40 nM pUC19 was used in the presence of 20, 40, 80 and 160 nM LmuABC. The reaction was conducted in the reaction buffer (RB) with the same protocol for stopping the reaction and loading on the gel. For the protein component requirement for LmuABC catalyzed DNA degradation experiment (**Extended Data Fig. 12b)**, 40 nM of pUC19 was used with 100nM LmuBC (Lane 2), 100nM LmuA (heparin fraction) and 1mM ATP (Lane3), 100nM LmuBC and 100nM LmuA (heparin fraction) with 1mM ATP (Lane 4), 100nM LmuBC and 100nM LmuA (heparin fraction) without ATP (Lane 5), 100 nM LmuA_K57A_ (heparin fraction) with 1mM ATP (Lane 6), 100 nM LmuBC and 100 nM LmuA_K57A_ (heparin fraction with 1mM ATP (Lane 7). All reactions were carried out with the same reaction buffer and reaction stopping conditions.

### Native mass spectrometry (nMS) analysis of complexes

LmuABC and LmuABC-DNA bound samples were buffer-exchanged into 300 mM ammonium acetate pH 7.5, 0.01% Tween-20 using Zeba microspin desalting columns with a 7-kDa molecular weight cutoff (Thermo Scientific). A 2 - 3 µL aliquot of the sample was then loaded into a gold-coated quartz capillary tip that was prepared in-house and then electrosprayed into an Exactive Plus with extended mass range (EMR) instrument (Thermo Fisher Scientific) with a static direct infusion nanospray source ^65^. The typical MS parameters for all samples included: spray voltage, 1.20 - 1.22 kV; capillary temperature, 150 °C; in-source dissociation, 10 V; S-lens RF level, 200; resolving power, 8,750 or 17,500 at *m/z* of 200; AGC target, 1 × 10^6^; maximum injection time, 200 ms; number of microscans, 5; total number of scans, at least 100. Additional nMS parameters were injection flatapole, 8 V; interflatapole, 7 V; bent flatapole, 4 - 6 V; high energy collision dissociation (HCD), 150 - 200 V; ultrahigh vacuum pressure, 3.4 - 4.8 × 10^−10^ mbar. Mass calibration in positive EMR mode was performed using cesium iodide. For data processing, the collected nMS spectra were visualized using Thermo Xcalibur Qual Browser (v. 4.2.47). Spectral deconvolution was performed using UniDec v. 4.2.0 ^66,67^ with the following general parameters: for background subtraction (if applied), subtract curve 10; smooth charge state distribution, enabled; peak shape function, Gaussian; Degree of Softmax distribution (beta parameter): 10 - 20. The expected masses for the component proteins include LmuA: 44,615.3 Da, LmuB: 74,619.0 Da, and LmuC: 20,286.8 Da. The observed mass deviations (calculated as the % difference between the measured and expected masses relative to the expected mass) ranged from 0.004 - 0.03%. The generation of LmuABC-dsDNA complex was undertaken with a 20-mer dsDNA duplex (5’-GTGATAGTTAGAAACGTAAT-3’ sequence and its complementary strand, expected mass : 12,230.1 Da).

## Data and code availability

All standardized datasets are available as follows: cryo-EM maps have been deposited in the Electron Microscopy Data Bank (EMDB) under the accession codes EMDB: EMD-49911 (LmuA_conformation 1), EMD-49915 (LmuA_conformation 2_asymmetric), EMD-49922 (LmuABC_apo), EMD-49934 (LmuABC-DNA). The corresponding atomic coordinates of the cryo-EM structures have been deposited in the Protein Data Bank (PDB) under the accession codes PDB: 9NXX (LmuA_conformation 1), PDB: 9NY1 (LmuA_conformation 2_asymmetric), PDB: 9NY5 (LmuABC_apo) and PDB: 9NYG (LmuABC-DNA).

Alignements, phylogenetic trees, DefenseFinder models and profiles, AF3 models, Lambda-vir genome and vector sequences are available in Zenodo at https://doi.org/10.5281/zenodo.15120681.

## Authors contributions

SHS, DJP and AB conceived the study. AC and ZZ undertook biochemical and cryo-EM studies under the supervision of DJP. MJdlC assisted with cryo-EM data processing and data collection. PDBO performed nMS analysis in the lab of BTC. MH, RL and MAA performed *in vivo* experiments, plaque assay, designed and built catalytic and interface mutants and toxicity assays. MH and AHdV undertook structural modeling with AF3, models analysis, comparisons and representation. MH performed comparative genomics, phylogenies, functional annotation. All authors participated in writing and editing the manuscript.

## Acknowledgements

The MDM lab is grateful to its members and Enzo Poirier for their useful comments on earlier versions of the manuscript. We thank F.T. Hoffmann and G.D. Lampe for guidance and technical support on ChIP-seq experiments and data analysis. Several bioinformatic analyses were performed on the Core Cluster of the Institut Français de Bioinformatique (IFB) (ANR-11-INBS-0013).

