## Extended Data Table 1 for "Structural basis for Lamassu-based antiviral immunity and its evolution from DNA repair machinery"

**Cryo-EM data collection, refinement and validation statistics**

|  | LmuA_symmetric tetramer  (EMD-49911)  (PDB:9NXX) | LmuA_asymmetric tetramer  (EMD-49915)  (PDB:9NY1) | LmuABC_apo  complex  (EMD-49922)  (PDB:9NY5) | LmuABC_DNA complex  (EMD-49934)  (PDB:9NYG) |
| --- | --- | --- | --- | --- |
| **Data collection and processing** |  |  |  |  |
| Magnification | 22,500 | 22,500 | 165,000 | 165,000 |
| Voltage (kV) | 300 | 300 | 300 | 300 |
| Electron exposure (e–/Å^2^) | 53 | 53 | 29.28 | 29.28 |
| Defocus range (μm) | -0.8 to -2.2 | -0.8 to -2.2 | -0.8 to -2.2 | -0.8 to -2.2 |
| Pixel size (Å) | 1.064 | 1.064 | 0.725 | 0.725 |
| Symmetry imposed | C1 | C1 | C1 | C1 |
| Initial particle images (no.) | 7,002 | 7,002 | 11,653 | 11,653 |
| Final particle images (no.) | 6,559 | 6,559 | 11,653 | 11,653 |
| Map resolution (Å)  FSC threshold | 3.13  0.143 | 3.40  0.143 | 3.21  0.143 | 2.93  0.143 |
| Map resolution range (Å) | 3.0 to 5.0 | 3.0 to 5.0 | 3.0 to 5.5 | 2.8 to 5.5 |
| **Refinement** |  |  |  |  |
| Initial model used (PDB code) | AF3 | AF3 | AF3 | AF3 |
| Map sharpening *B* factor (Å^2^) | -129.9 | -125.4 | -87.3 | -62.6 |
| Model composition  Non-hydrogen atoms  Protein residues  Ligands | 11,911  1,465  0 | 11,518  1412  0 | 6,747  838  0 | 10,823  1,255  0 |
| *B* factors (Å^2^)  Protein  Ligand | 164.50  NA | 195.25  NA | 162.74  NA | 173.32  NA |
| Nucleotide | NA | NA | NA | 188.98 |
| R.m.s. deviations  Bond lengths (Å)  Bond angles (°) | 0.030  1.028 | 0.034  1.544 | 0.004  0.898 | 0.008  1.221 |
| Validation  MolProbity score  Clashscore  Poor rotamers (%) | 2.17  7.78  2.66 | 1.93  6.98  1.18 | 1.83  6.20  1.46 | 2.44  7.59  6.26 |
| Ramachandran plot  Favored (%)  Allowed (%)  Disallowed (%) | 93.60  5.92  0.48 | 91.76  8.17  0.07 | 94.63  5.37  0.00 | 93.73  6.10  0.16 |
