## Extended Data Figures 1-17 for "Structural basis for Lamassu-based antiviral immunity and its evolution from DNA repair machinery"

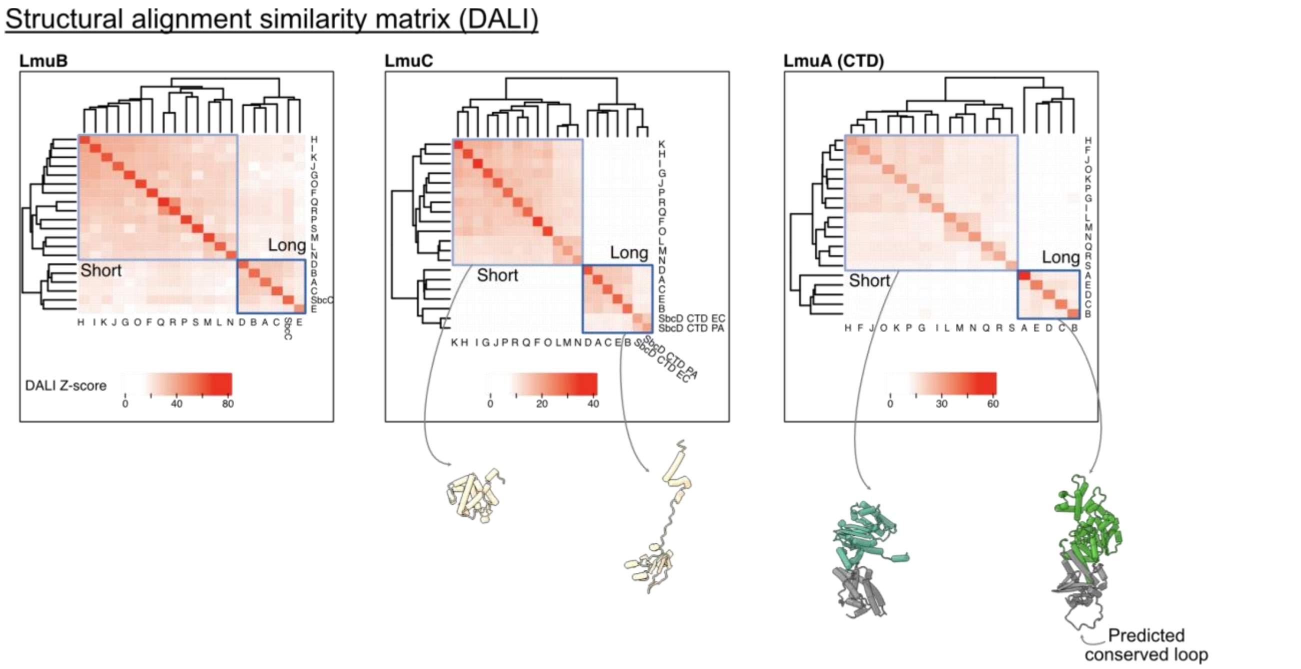


**Extended Data Fig. 1. 3D structure comparisons of Alphafold 3 models with DALI webserver**. The distance matrices were clustered by hierarchical clustering, and the dendrograms are shown on top of the plots. Proteins associated with short and long Lamassus form separate clusters. Representative monomers of LmuC and LmuA are shown below; For LmuA, the CTD is coloured in grey, and the effector in green.

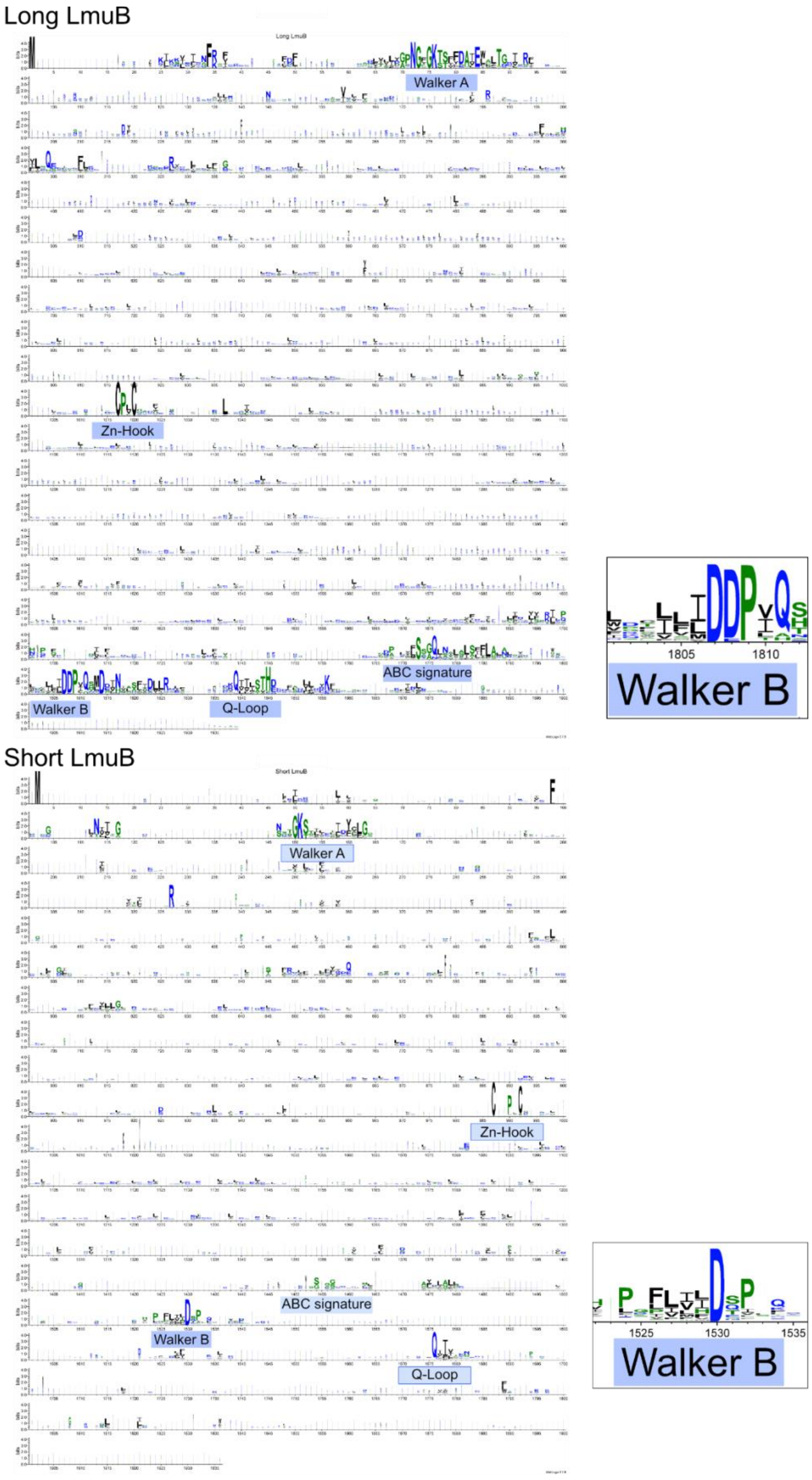


**Extended Data Fig. 2**. **Alignments of short and long LmuB.**

WebLogo of Long (upper) and Short (lower) LmuB based on the alignment of representative sequences in Figure 1. Motifs are annotated below the consensus sequence. Walker B consensus zoom-in are represented on the right.

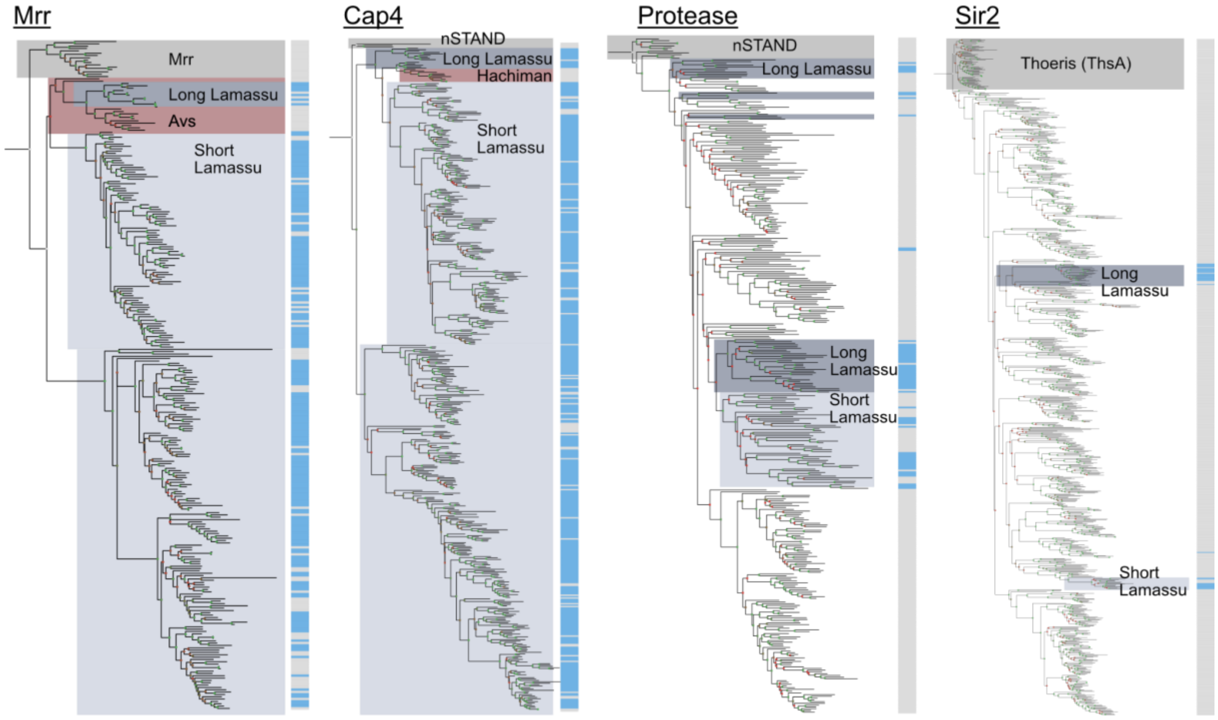


**Extended Data Fig. 3. Phylogenetic analyses of the Mrr, Cap4, Protease, and Sir2 effector domains**.

Sequences from representative Lamassu systems (from Figure 1) were aligned using MAFFT (Linsi), and the N-terminal effector domain sequences were extracted for further analysis. Profile HMMs were constructed from each alignment and used to search against the RefSeq database with default parameters and an E-value threshold of <1e-5. The retrieved sequences were clustered using MMSeqs2 at 60% identity and coverage, aligned with MAFFT (--auto) alongside distant homologs for rooting, and further processed to extract the effector domain, ensuring the removal of any additional domains. Phylogenetic trees were inferred using IQ-TREE2 (-B 1000, -mset WAG,Q.pfam) and visualized with TreeViewer. Rooting sequences included the Mrr restriction enzyme (Mrr domain), Avs-Cap4 from the nSTAND family (Cap4 domain), Avs-Protease from the nSTAND family (Protease domain), and ThsA from the Thoeris system (Sir2 domain). Clades were annotated based on their associated defense system, as identified by DefenseFinder, with blue bars marking sequences belonging to Lamassu systems. UFB is represented with internal node squares, <80 (red) and >80 (green).

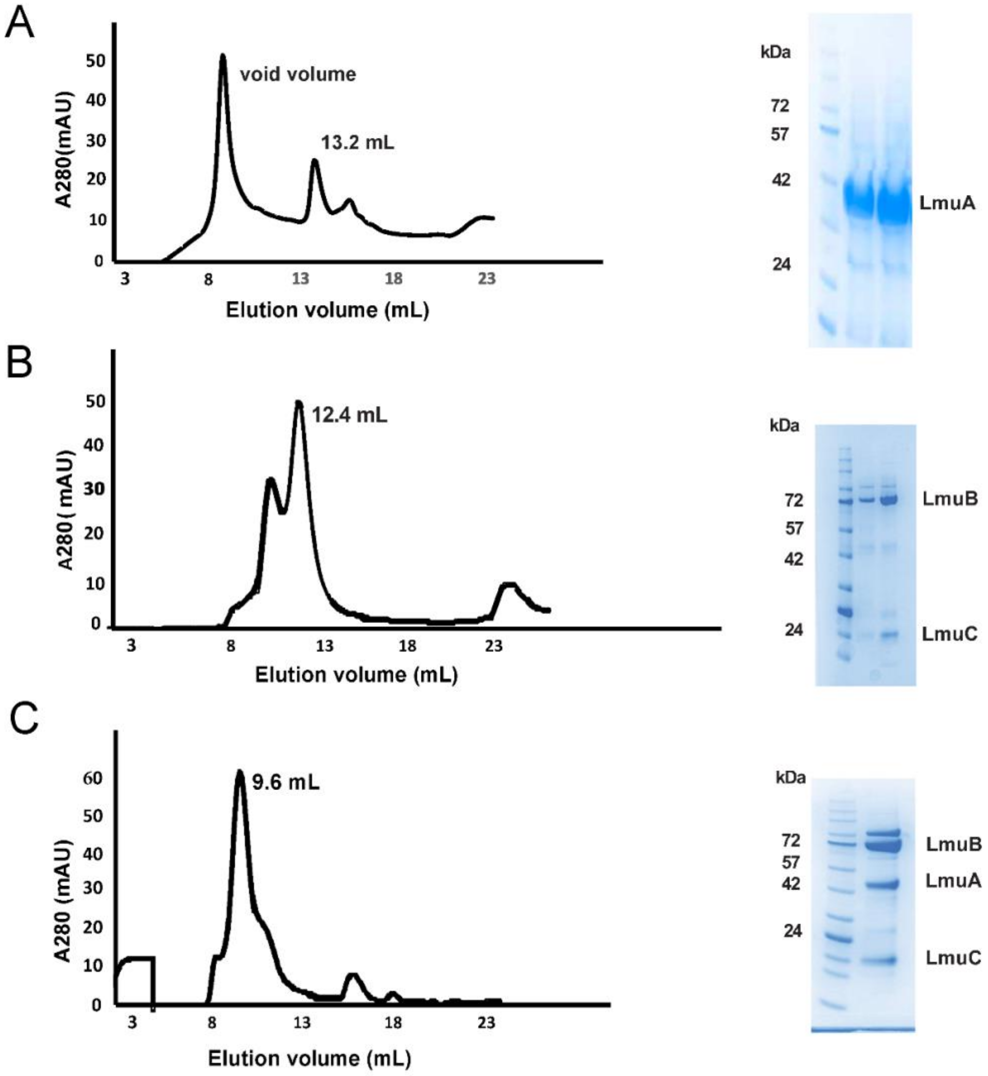


**Extended Data Fig. 4. Lamassu protein purification.**

**(a)** SEC profile (superdex 200 10/300 gl, Cytiva) and SDS-PAGE gel of LmuA indicating aggregation in the column. The SDS-PAGE gel stained with Coomassie brilliant blue, indicates protein peak fraction concentrated from the void volume in lane1 indicating severe aggregation. Concentrated protein fraction from 13.2 mL peak had a tendency to precipitate upon concentration. **(b)** SEC profile (superdex 200 10/300 gl, cytiva) and SDS-PAGE gel of LmuBC indicating complex formation. The SDS-PAGE gel stained with Coomassie brilliant blue indicates two fractions from the peak fraction of 12.4 mL peak. **(c)** SEC profile (superdex 200 10/300 gl, Cytiva) and SDS-PAGE gel of LmuABC indicating complex formation after mixing separately expressed LmuA and LmuBC. The SDS-PAGE gel stained with Coomassie brilliant blue indicates peak fraction at 9.6 mL.

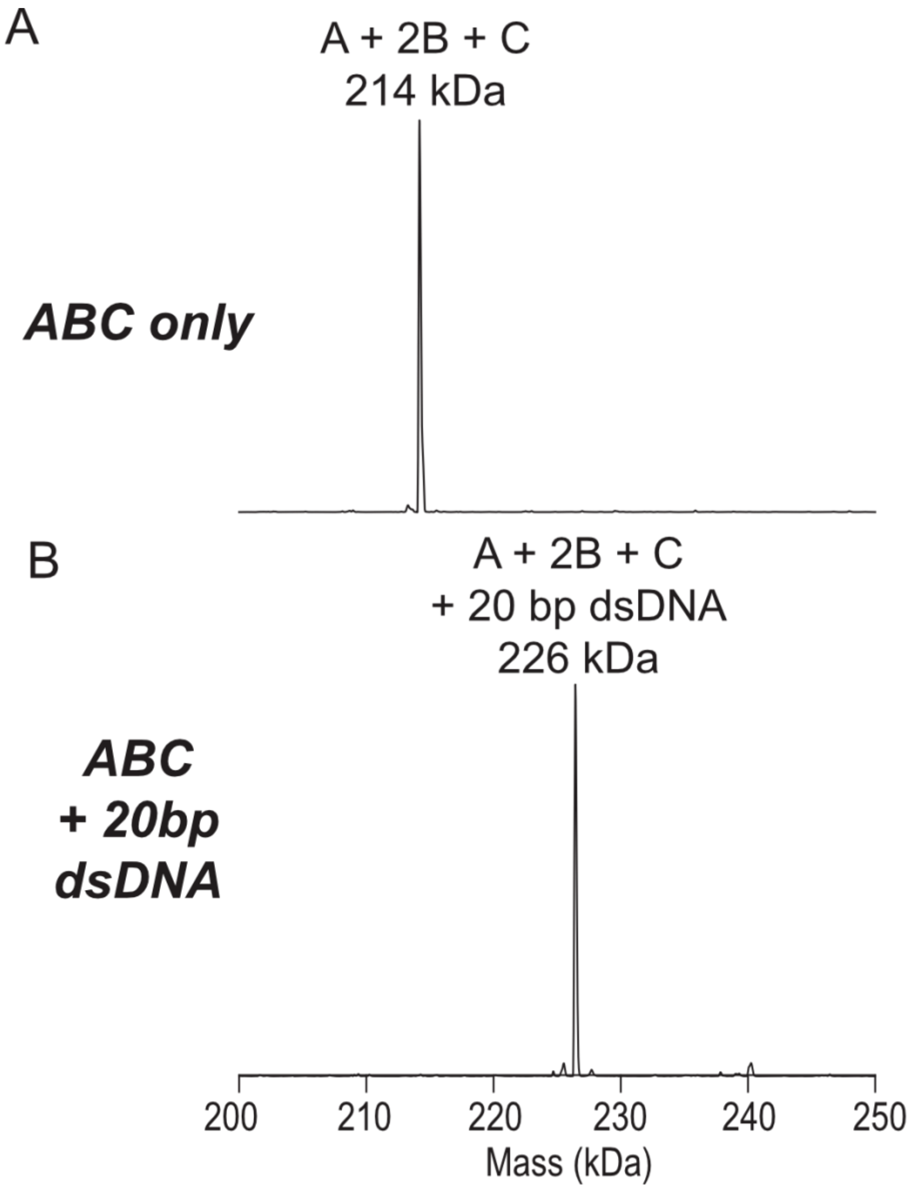


**Extended Data Fig. 5. Native mass spectrometry (nMS) analysis of the LmuABC complex in the apo- and 20-bp DNA-bound states.**

Deconvolved nMS spectra for the (**a**) apo LmuABC and (**b**) LmuABC incubated with 20-bp dsDNA. nMS measurements establish that the complex has a stoichiometry of LmuA$_1$B$_2$C$_1$ and binds one 20-bp dsDNA. The expected masses for the components are LmuA: 44,615 Da, 2LmuB: 149,238 Da, LmuC: 20,287 Da and 20-bp dsDNA: 12,230 Da. The predicted masses from the established subunit stoichiometry are LmuA$_1$B$_2$C$_1$: 214,140 Da and LmuA$_1$B$_2$C$_1$ + 20 bp dsDNA: 226,371 Da.

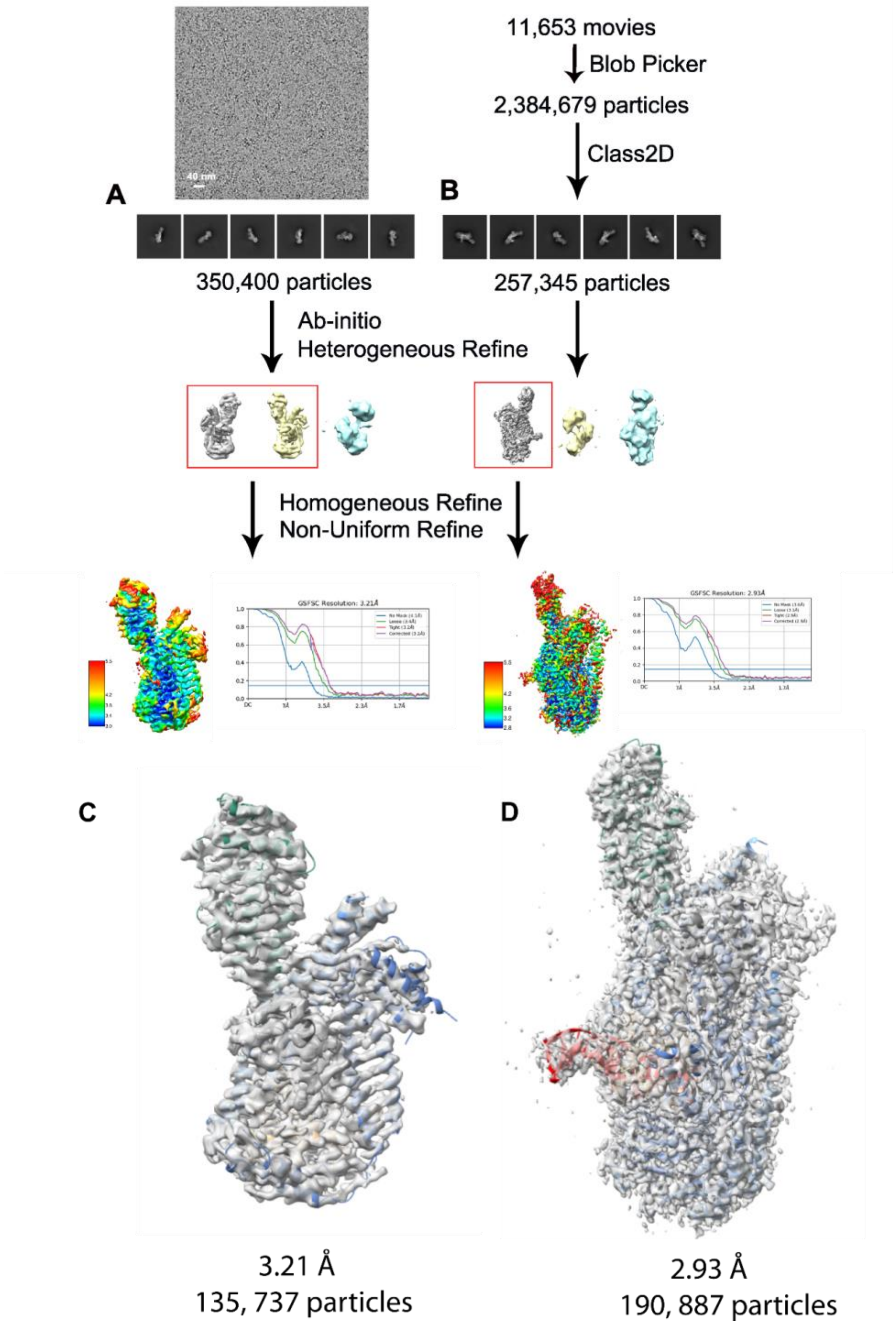


**Extended Data Fig. 6. Cryo-EM work-flow for LmuA$_1$(K57A)B$_2$C$_1$ complex bound to 20-bp DNA structure determination.**

**(a)** Cryo-EM data processing work-flow and statistics of data set processing for apo-LmuABC component. **(b)** Cryo-EM data processing work-flow and statistics of data set processing for one LmuABC bound to dsDNA component. **(c)** Cryo-EM model and map of apo-LmuABC. **(d)** Cryo-EM model and map of LmuABC-DNA complex.

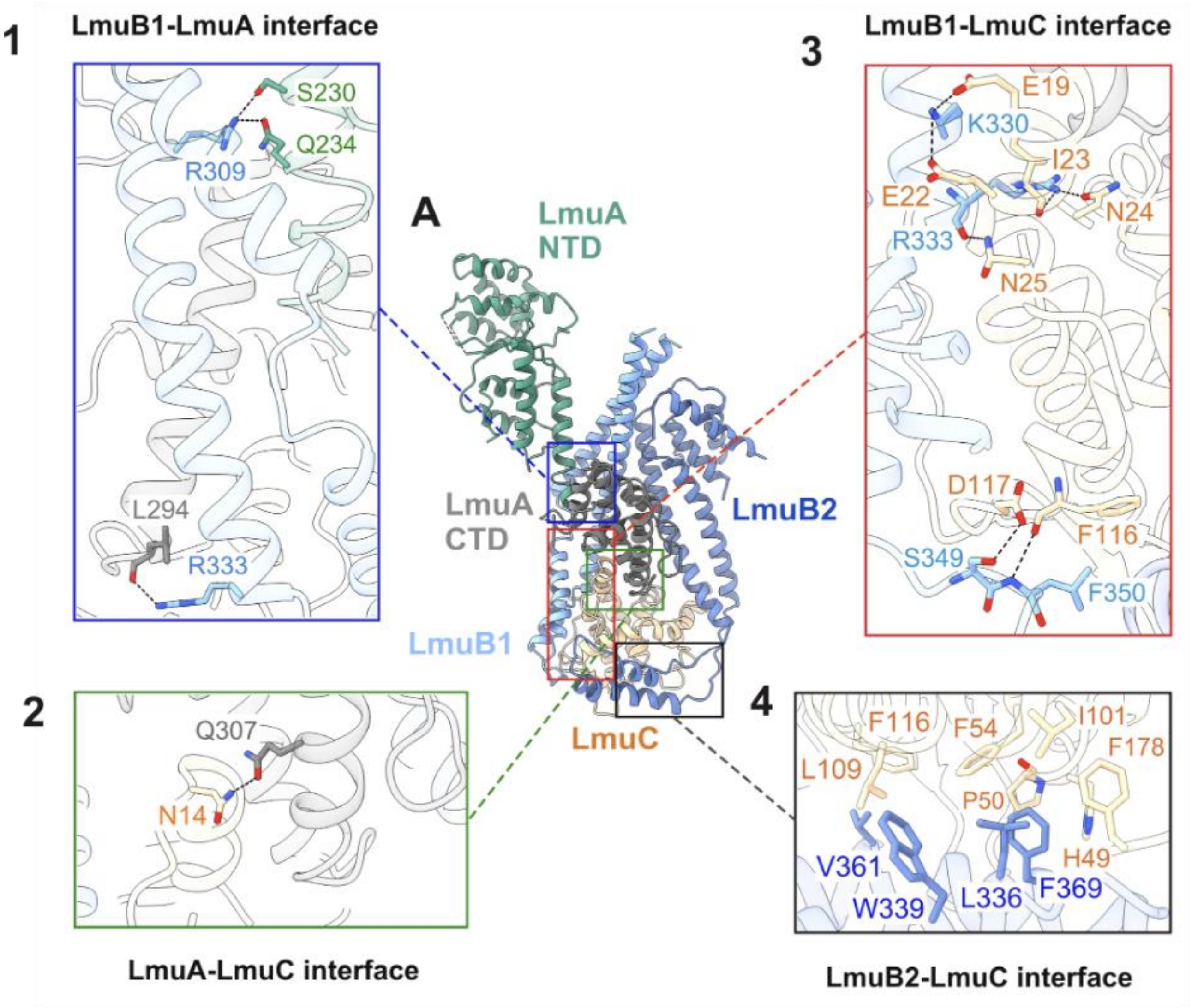


**Extended Data Fig. 7. Structure of apo-LmuABC.**

The LmuABC structure is composed of $LmuA_1B_2C_1$. We observe a core topology formed by head-distal coiled coils of the pair of LmuBs (in light and dark blue), the LmuC (beige) and the C-terminal domain of LmuA (grey) **(1-4)** Details of hydrogen bonding (panels 1, 2, 3) and hydrophobic interactions (panel 4) in the LmuABC structure are shown in expanded box segments labeled 1 to 4.

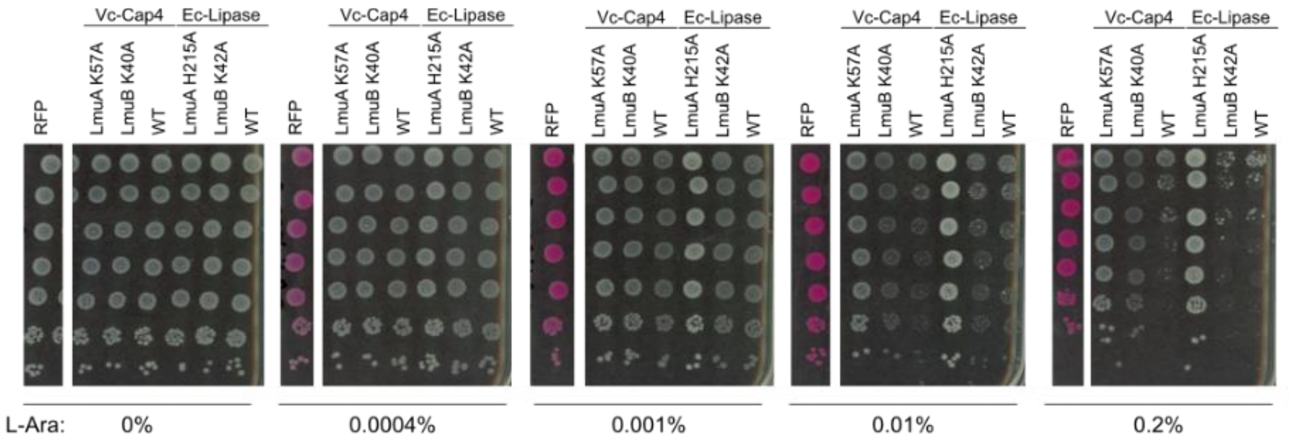


**Extended Data Fig. 8: Toxicity of Lamassu at various concentrations of inducer.**
Overnight cultures of *E. coli* DH10B transformed with a pBAD plasmid encoding WT and associated catalytic mutants of LmuB (Walker A catalytic K→A) and LmuA (K57A or H215A) were cultured with Carbenicillin (100 uM) and Glucose (0.01%). The addition of glucose is necessary to avoid leaky expression of the system which leads to the accumulation of spontaneous mutants. Cultures were then serially diluted in LB and plated on increasing amounts of L-Arabinose (L-Ara). Toxicity is apparent starting from 0.01% L-Arabinose, except for LmuA catalytic mutants. DH10B+pBAD-RFP is shown as control.

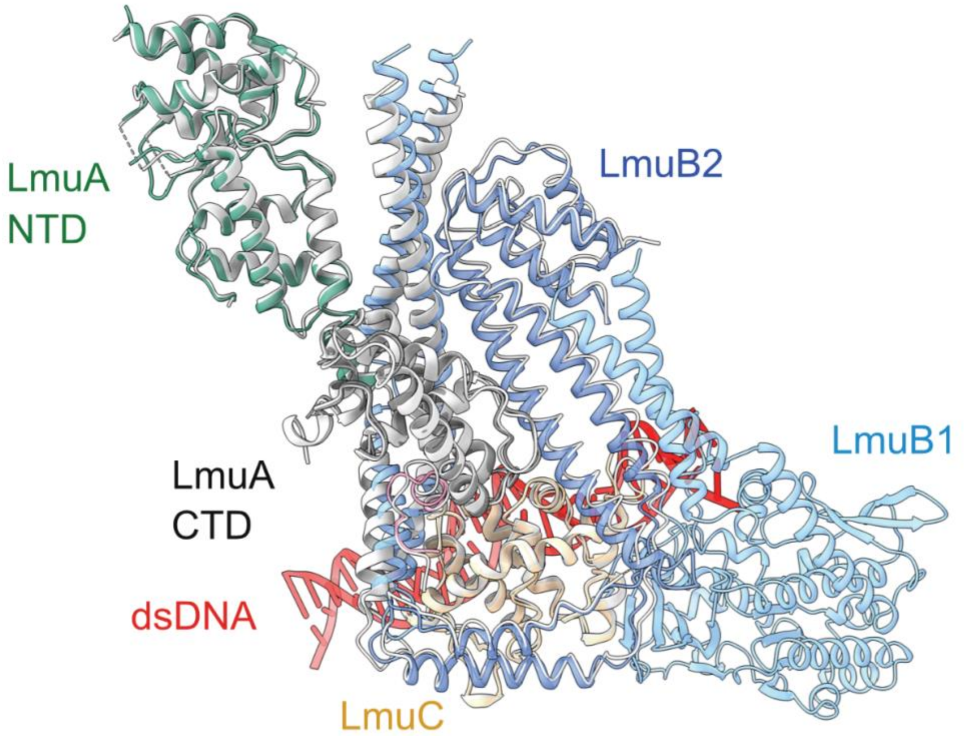


**Extended Data Fig. 9**. **Comparison of Apo- and dsDNA-bound complexes.**
Superposition of cryo-EM structures of apo-$LmuA_1B_2C_1$ (in silver) and $LmuA_1B_2C_1$-DNA (in color) with an RMSD of 0.795.

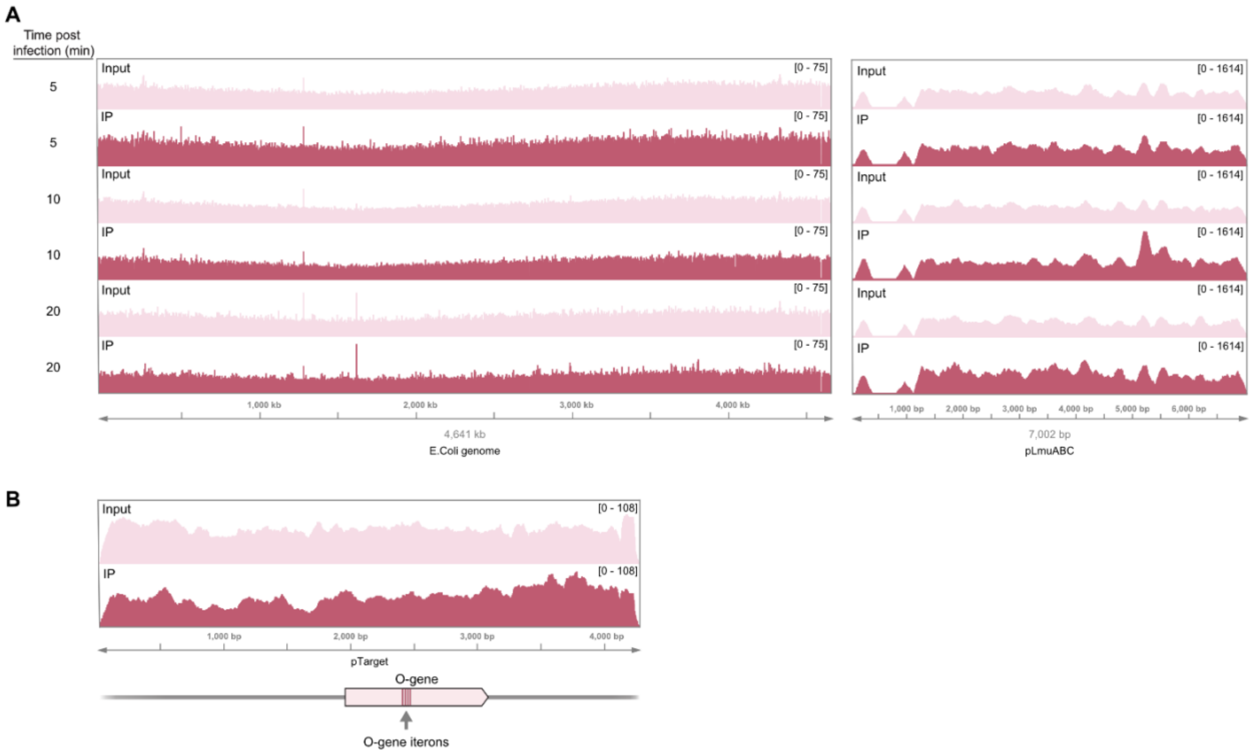


**Extended Data Fig. 10. Analysis of additional ChIP-seq data.**

**(a)** ChIP-seq data from experiments shown in **Figure 3D** were mapped to the *E. coli* genome (left) and Lamassu expression plasmid (right), revealing an absence of enriched peaks. The genome- and plasmid-wide graphs show normalized read coverage at various time points after Lambda-vir infection, with input and IP samples colored in pink and maroon, respectively. **(b)** ChIP-seq experiments were repeated in the absence of Lambda-vir, but in the presence of an accessory plasmid containing an ectopically encoded full length O-gene sequence. No LmuB enrichment was observed at the O binding repeat iterons, unlike experiments with Lambda-vir, suggesting that Lamassu specifically recognizes these sequences in the context of phage replication.

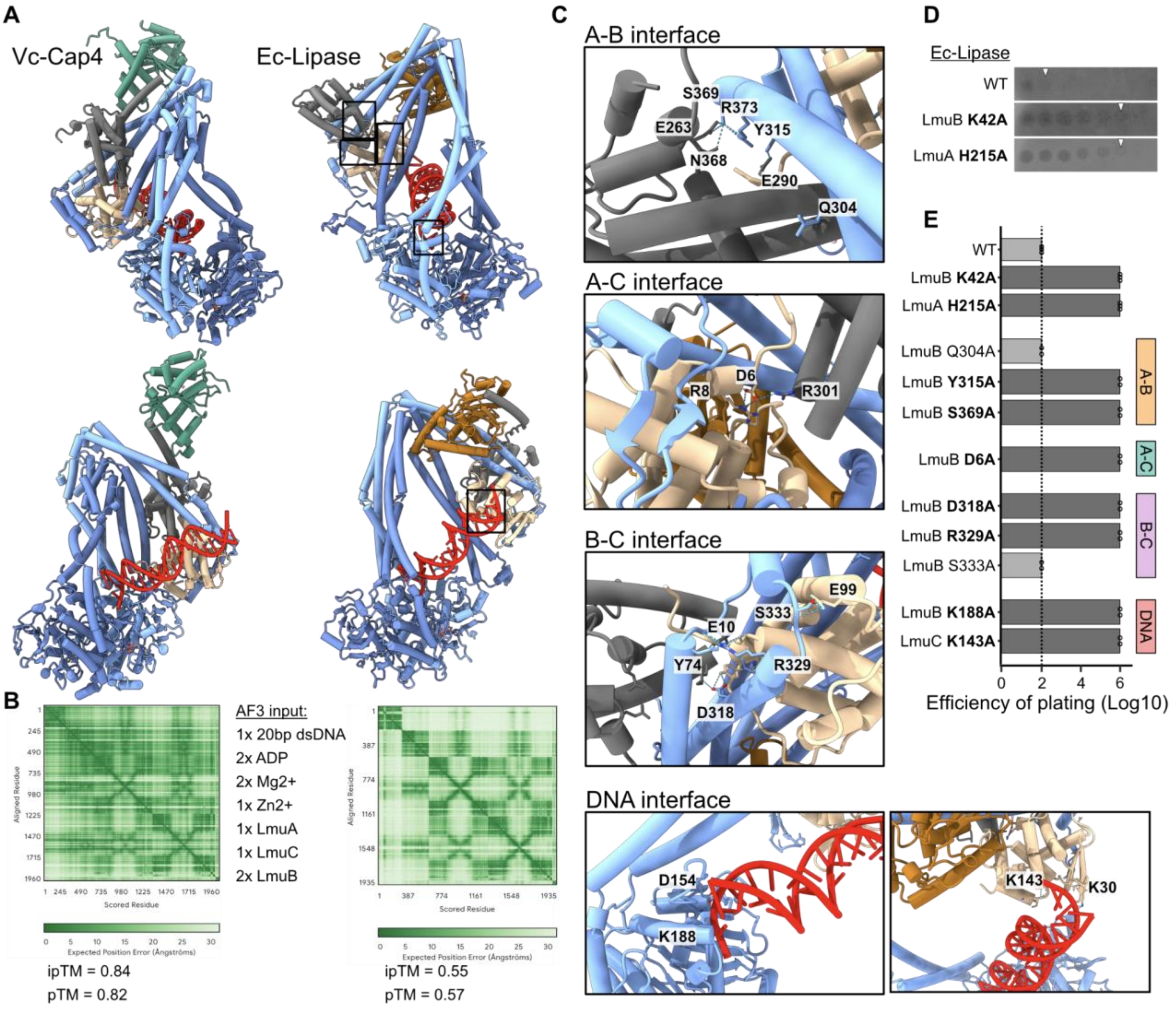


**Extended Data Fig. 11**. **Comparison of Vc-Cap4 and Ec-Lipase AF3 multimer models.**
**(a)** AF3 models of Vc-Cap4 and Ec-Lipase with the stoichiometry learned from nMS and Cryo-EM of Vc-Cap4. Whole models comparisons. **(b)** The predicted alignment error (PAE) matrix are shown together with the input given to AF3 and resulting iPTM and pTM scores. Zn2+ was added since the Ec-Lipase system has a CxxC motif. **(c)** Zoom-in panels of the Ec-Lipase model showing residue interactions at distinct interfaces. **(d)** Experimental validation of Ec-Lipase. Plaque assay against Lambda-vir, 10-fold dilutions from left to right. Ec-Lipase was expressed from a plasmid under pBAD, with 0.001% L-arabinose. LmuB K42A is a mutant of the catalytic Lysine of the Walker A motif. LmuA H215A is a mutant of the catalytic histidine of the catalytic triad of lipases. **(e)** Key residues involved in side chain interactions at interfaces. Interfaces are annotated on the right. Efficiency of plating is the maximum 10-fold dilution factor at which lysis is observed.

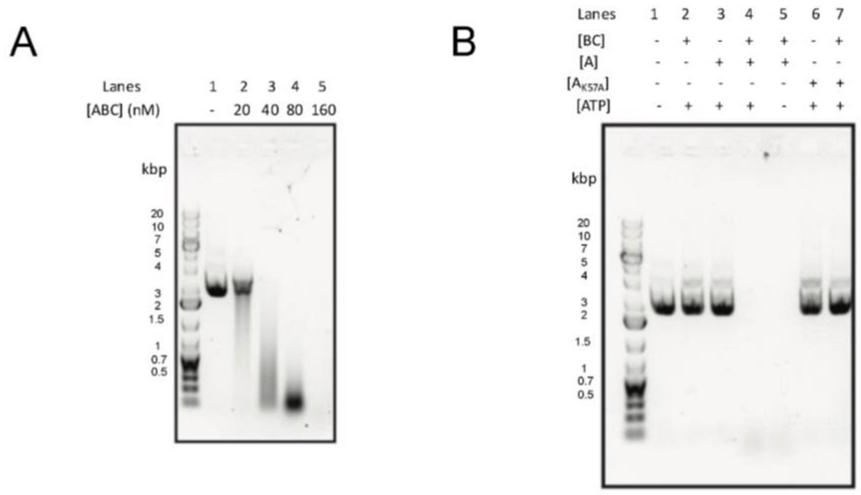


**Extended Data Fig. 12. Biochemical assays at high LmuABC concentrations.**
(**a**) Concentration dependency experiment showcasing the degradation of pUC19 at increasing LmuABC concentrations ranging from 20nM to 160nM. For all reactions 40 nM pUC 19 was used. (**b**) pUC19 degradation in the presence of various Lamassu protein components. 40 nM pUC19 and 100nM of each protein (LmuBC, $LmuA_{K57A}$, LmuA) with 1mM ATP was used for their respective reactions.

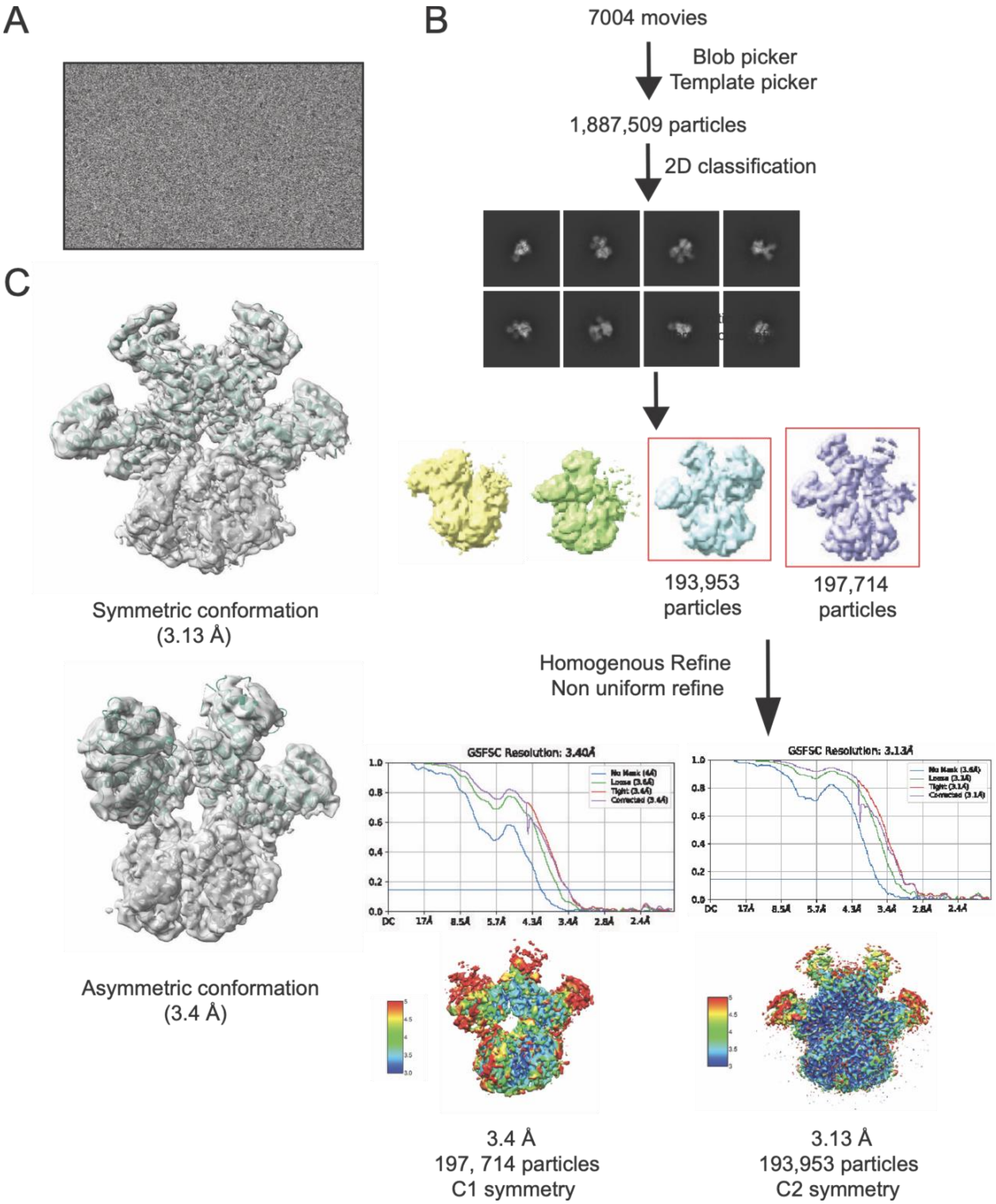


**Extended Data Fig. 13**. **Cryo-EM work-flow for LmuA structure determination.**

**(a)** Cryo-EM raw micrograph of the LmuA tetramer. **(b)** Cryo-EM work flow and statistics of the LmuA dataset processing. **(c)** Cryo-EM model and map of symmetric (top) and asymmetric (bottom) LmuA tetrameric conformations.

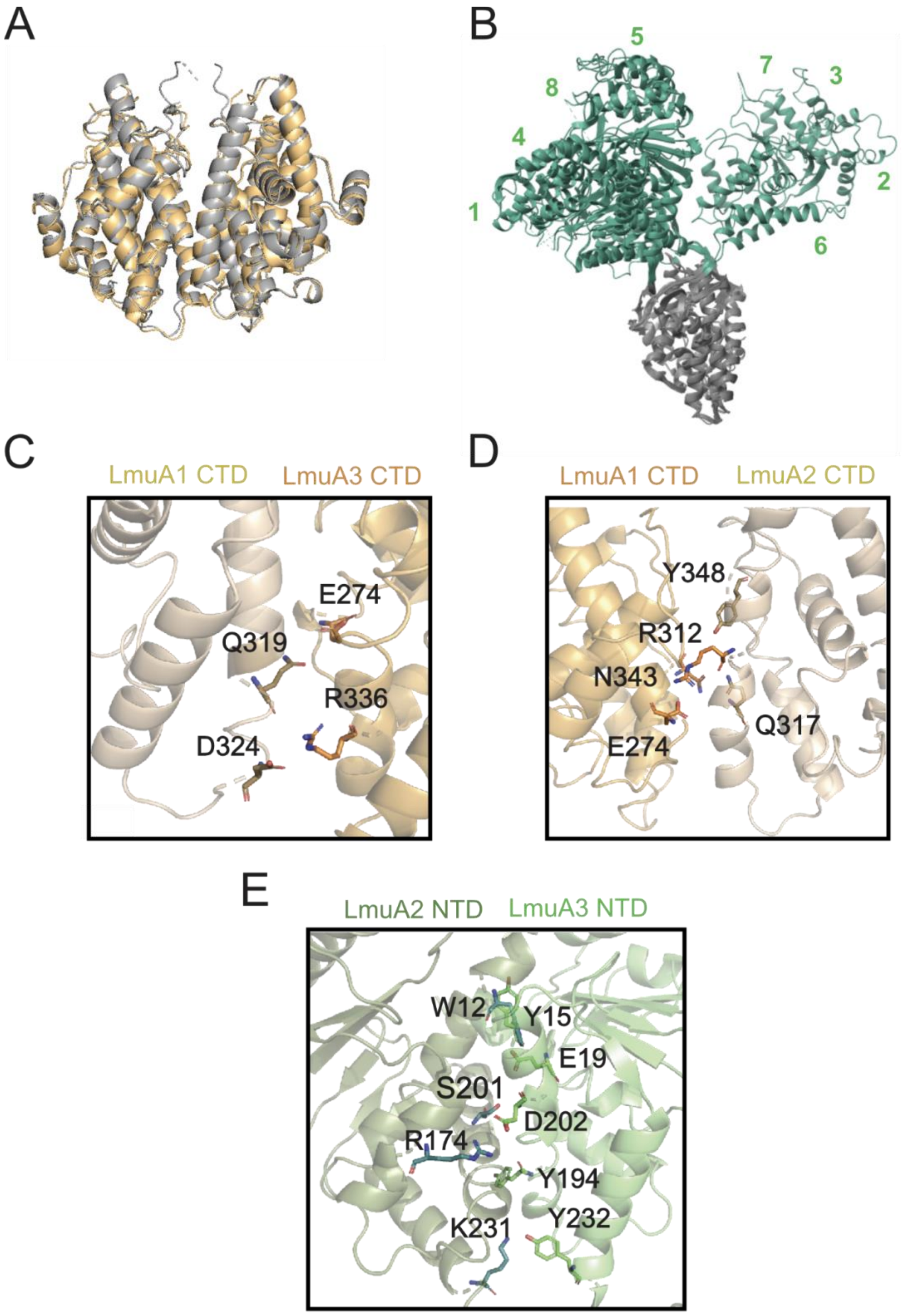


**Extended Data Fig. 14**. **Comparison of asymmetrical and symmetrical tetramer cryo-EM structures.**

**(a)** Superposition of CTD segments of symmetric (in silver) and asymmetric (in color) conformations of the LmuA tetramer. **(b)** Alignment of monomers of the symmetric (labeled 1 to 4) and asymmetric (labeled 5 to 8) structures of LmuA tetramer, following superposition of their CTD domains. **(c,d)** Interfacial interactions associated with the CTDs of the symmetrical LmuA tetramer within interface A between monomers labeled 1 and 3 (panel C) and in interface B between monomers labeled 1 and 2 (panel D). **(e)** Extensive interactions between four-helix bundles of symmetry-related N-terminal domains in the symmetric LmuA tetramer for monomers labeled 2 and 3.

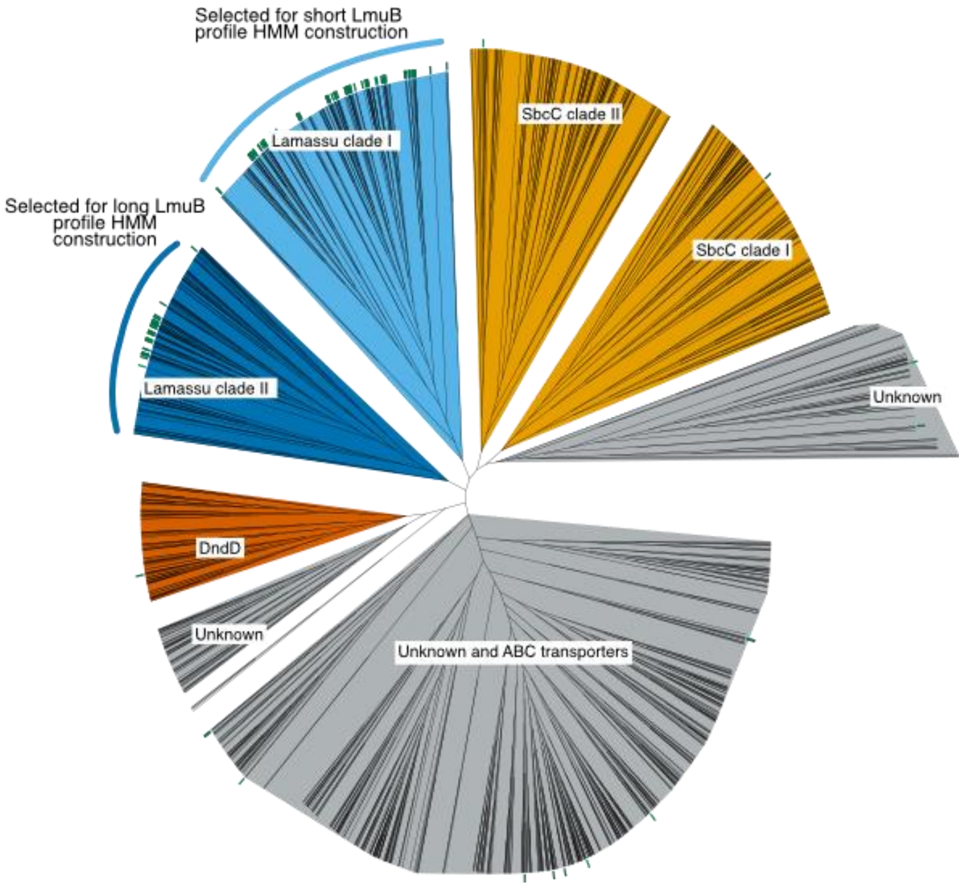


**Extended Data Fig. 15. Phylogeny of LmuB homologs.**
Unrooted phylogenetic tree of the LmuB profile HMM hits. Green rectangles correspond to gene families with neighbors annotated as LmuA or LmuC by a previous version of DefenseFinder and ABC-3C PFAM profiles.

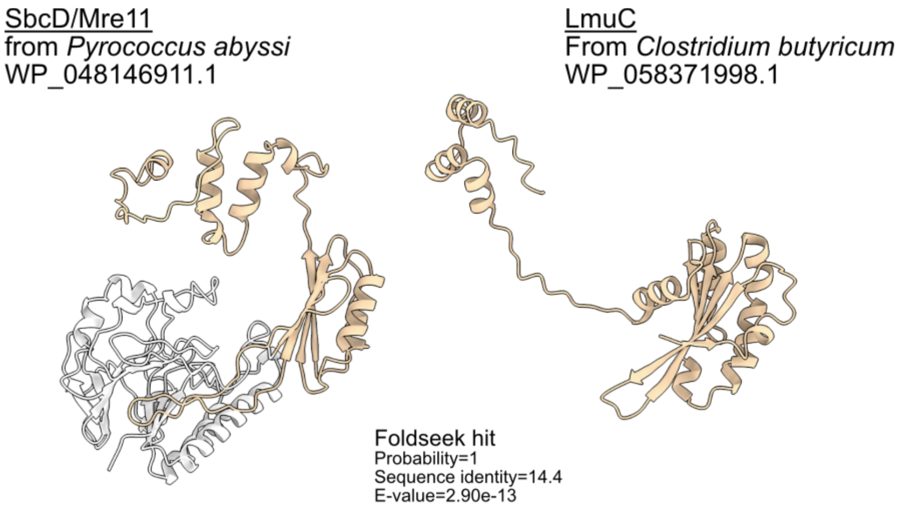


**Extended Data Fig. 16. Structural comparison of SbcD to long LmuC.**
Best hit of the long LmuC from *Clostridium butyricum* in the AFDB-Swissprot Database identified with Foldseek (right panel). Match protein is SbcD/Mre11 from the genome of the archaeon *Pyrococcus abyssi* (left panel). Domains were matched through the SbcC binding domain of SbcD (CTD) to the whole LmuC protein (highlighted in beige).

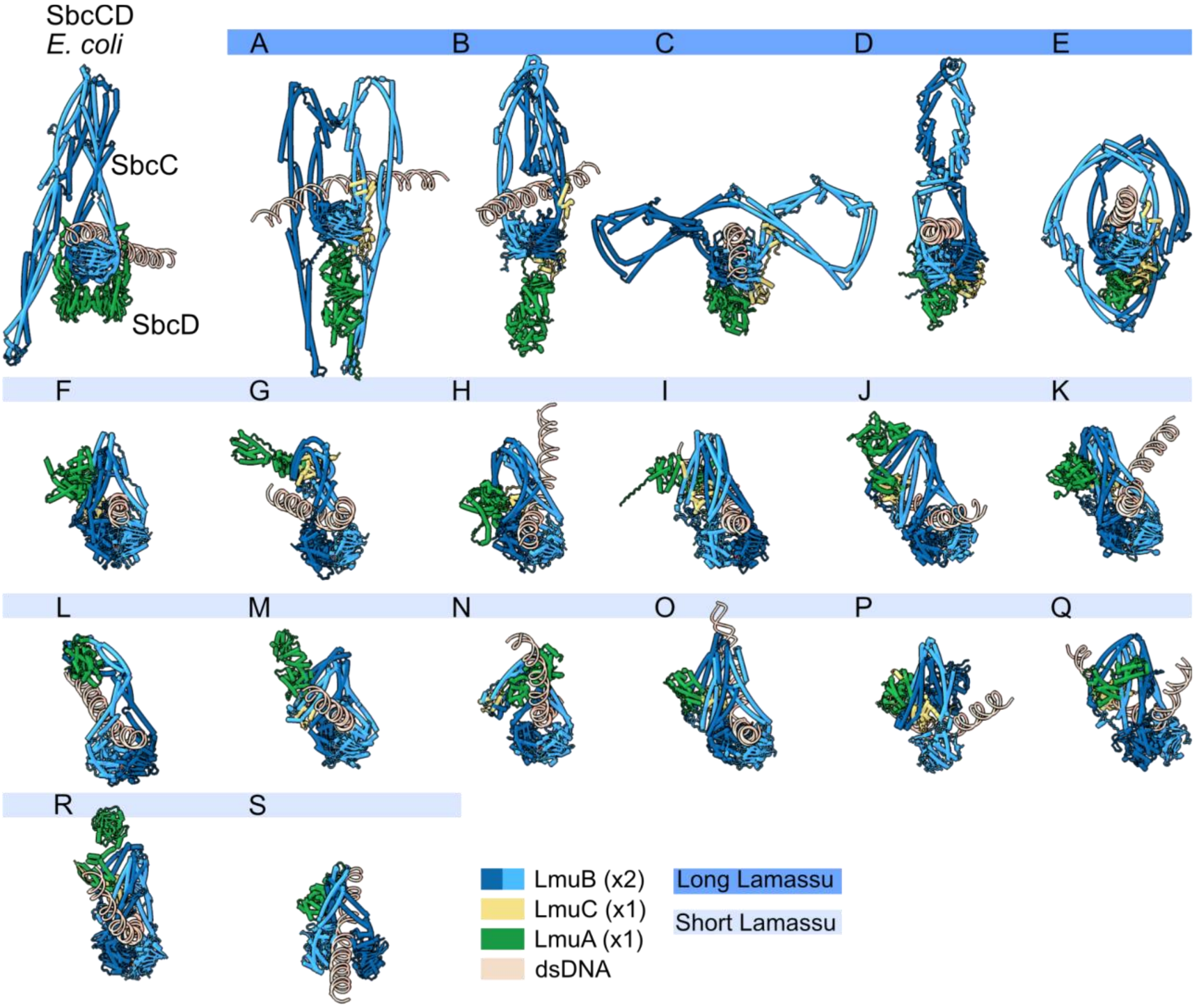


**Extended Data Fig. 17. AlphaFold3 models of SbcCD, long and short Lamassus.**
Multimer AF3 models of SbcCD and the 19 identified clades of Lamassu with fixed stoichiometry (1x LmuA, 1x LmuC, 2x LmuB), co-factors (2x $Mg^{2+}$, 1x $Zn^{2+}$, 2x ADP) and 50 bp dsDNA sequence. Long Lamassu (A-E) and Short Lamassu (F-S). SbcCD is colored accordingly, SbcC (blue) and SbcD (green). The predicted molecular complexes are similar within, but different between, long and short Lamassu.
